# Defining the networks that connect RNase III and RNase J-mediated regulation of primary and specialized metabolism in *Streptomyces venezuelae*

**DOI:** 10.1101/2025.01.15.633244

**Authors:** Meghan A.D. Pepler, Emma L. Mulholland, Freddie R. Montague, Marie A. Elliot

## Abstract

RNA metabolism involves coordinating RNA synthesis with RNA processing and degradation. Ribonucleases play fundamental roles within the cell, contributing to the cleavage, modification, and degradation of RNA molecules, with these actions ensuring appropriate gene regulation and cellular homeostasis. Here, we employed RNA-sequencing to explore the impact of RNase III and RNase J on the transcriptome of *Streptomyces venezuelae*. Differential expression analysis comparing wild type and RNase mutant strains at distinct developmental stages revealed significant changes in transcript abundance, particularly in pathways related to multicellular development, nutrient acquisition, and specialized metabolism. Both RNase mutants exhibited dysregulation of the BldD regulon, including altered expression of many cyclic-di-GMP-associated enzymes. We also observed precocious chloramphenicol production in these RNase mutants and found that in the RNase III mutant, this was associated with PhoP-mediated regulation. We further found that RNase III directly targeted members of the PhoP regulon, suggesting a link between RNA metabolism and a regulator that bridges primary and specialized metabolism. We connected RNase J function with translation through the observation that RNase J directly targets multiple ribosomal protein transcripts for degradation. These findings establish distinct, but complementary roles for RNase III and RNase J in coordinating the gene expression dynamics critical for *S. venezuelae* development and specialized metabolism.

## INTRODUCTION

Post-transcriptional regulation is a critical component of cellular control in bacteria. The constant turnover of RNA can facilitate rapid changes in gene expression and the stability of a given transcript is largely governed by its accessibility to ribonucleases (RNases) [1]. RNases can degrade RNA from within a transcript (endoribonucleases) or from a terminal end (exoribonuclease). In all bacteria, RNA turnover is primarily mediated by degradosome complexes [2–5]. The degradosome has been best studied in the Gram-negative bacterium *Escherichia coli,* where the complex is assembled around the large endoribonuclease RNase E [3, 6]. In this complex, RNase E associates with the exoribonuclease PNPase, the glycolytic enzyme enolase, and the DEAD-box RNA helicase RhlB. The degradosome in Gram-positive bacteria involves a different suite of RNases, as most Gram-positive bacteria do not encode RNase E homologs. In *Bacillus subtilis,* the membrane-anchored endoribonuclease RNase Y is the primary scaffold for the degradosome complex, with interacting proteins including a DEAD-box helicase, enolase, PNPase, and RNase J, which has dual 5ʹ-3ʹ exoribonuclease and endoribonuclease activity [7–10]. RNase E and RNase Y both preferentially cleave 5ʹ monophosphorylated RNA molecules [10]. Thus, RNA decay is typically initiated by a preliminary endonucleolytic cleavage event that generates monophosphorylated 5ʹ ends, or by a pyrophosphohydrolase (*e.g.,* RppH) that converts 5ʹ triphosphates to monophosphates [2]. In addition to a suite of diverse single-strand-specific RNases, most bacteria also encode the double-stranded RNA-specific RNase III [11]. This enzyme is highly conserved across bacteria and eukaryotes and its capacity to cleave double-stranded RNA makes it a critical regulator of both mRNAs bound by noncoding small RNAs (sRNAs) and highly structured RNAs like ribosomal RNAs (rRNAs) [11–14].

As investigations into RNase activity in bacteria have extended beyond the *E. coli* and *B. subtilis* model systems, it has become evident that there is diversity in the number and type of RNases employed by any given bacterium [3, 5, 7, 15–19]. The actinobacteria are a phylum of Gram-positive bacteria that includes *Streptomyces* and *Mycobacterium*. These bacteria encode a set of RNases that includes RNase III, RNase J, and contrary to what is observed in most other Gram-positive genera, RNase E [20]. *Streptomyces* have been studied extensively as a model of multicellular growth in bacteria and for their production of medically relevant specialized metabolites. The *Streptomyces* classical life cycle is defined by morphologically and metabolically distinct developmental stages [21]. The cycle begins with spore germination, which involves the emergence of one or two hyphal filaments that grow via tip extension and branching to form a dense, hyphal network. On solid media, reproductive growth initiates with the raising of aerial hyphae and culminates with the subdivision of the aerial cells into chains of unigenomic spores. In *Streptomyces* species that can differentiate in liquid medium (*e.g., Streptomyces venezuelae*), the growth stage that is analogous to aerial development is termed “fragmentation”, where the vegetative hyphae break apart before forming short chains of spores. It is during aerial development/fragmentation that *Streptomyces* typically initiate specialized metabolism. The cooperative regulation of multicellular development and specialized metabolism has been extensively studied in *Streptomyces,* but the contribution of RNases to the regulation of these processes has not been studied in detail.

We have previously observed that deleting *rnc* (RNase III) or *rnj* (RNase J) from *S. venezuelae* affects development, antibiotic production, normal sporulation, and ribosome assembly [22]. Loss of these enzymes leads to decreased production of the specialized metabolite jadomycin and the accumulation of 100S inactive ribosomes. Deleting *rnc* also causes the growing colony to peel away from solid medium, while the Δ*rnj* mutant develops more slowly in liquid medium than its wild type parent and is delayed in sporulation [22]. While the phenotypes of the *rnc* and *rnj* mutants are clear in *S. venezuelae*, the pathways by which their associated RNases exert their effects remain unknown. Here, we took a transcriptomic approach to understanding the changes in gene expression and transcript abundance over time within these two mutants. We observed that regulatory networks governing the progression of multicellular development, nitrogen assimilation, phosphate uptake, and specialized metabolism were significantly differentially expressed. The altered transcription profiles observed were similar for both mutants and were significantly different compared with their wild type parent strain; however, these parallel transcriptional changes yielded different phenotypic outcomes for the two mutant strains. We determined that RNase III, but not RNase J, likely contributes directly to the control of phosphate uptake in *S. venezuelae.* Our data also suggests that RNase III and RNase J affect the biogenesis of the translational machinery in distinct ways. Overall, our work emphasizes the complexity of RNase-based regulation and provides insights into how these enzymes affect the temporal regulation of development and specialized metabolism in *Streptomyces*.

## RESULTS

### RNA-sequencing analysis reveals a global impact of rnc and rnj deletion

To better understand how RNase III and RNase J affect *S. venezuelae* development and specialized metabolism, we undertook a transcriptomic approach. We isolated RNA from wild type and RNase mutant strains at distinct stages of classical development in liquid medium (vegetative growth, hyphal fragmentation, and – for the mutant strains – sporulation; we were unable to successfully extract intact RNA from sporulating wild type samples). We subjected the vegetative and hyphal fragmentation samples to differential expression analysis, comparing mutant and wild type strains. We also compared each mutant to itself over time, to assess trends in gene expression across developmental stages. For our differential expression analysis, we focussed our attention on genes that were reasonably well expressed (base mean >50) and that had significantly altered transcript abundance when comparing the wild type strain to either mutant (log_2_ fold change > |2|, with an adjusted p-value > 0.05).

When considering comparisons between wild type and RNase mutants during vegetative growth and hyphal fragmentation, we found 739 unique genes had significantly altered transcript levels in the Δ*rnc* mutant relative to the wild type, representing 9.8% of all annotated genes. In the Δ*rnj* mutant, 688 unique genes were significantly changed, representing 9.2% of all annotated genes. The Δ*rnc* and Δ*rnj* mutants shared 367 differentially expressed genes, which included 242 genes during vegetative growth, 171 genes during hyphal fragmentation, and 46 genes represented at both stages (**Figure 1A**). Intriguingly, we noted that these 367 shared genes tended to be similarly affected (either upregulated or downregulated in the same way, relative to wild type). In assessing the biological functions impacted by loss of these RNases, we sorted all significantly differentially expressed genes into functional categories defined by the Clusters of Orthologous Genes (COG) database [23]. For both the Δ*rnc* and Δ*rnj* mutants, among the COG categories that were most highly represented were those involved in transcription and signal transduction mechanisms (**Figure 1B**). This could, in part, explain the global shift in transcript abundance observed for these two mutants relative to their wild type parent, as altered levels of key transcriptional regulators could affect the transcript abundance of their associated regulons. We therefore sought to identify regulatory networks of interest within the signal transduction and transcription COG categories, where changing the abundance of a few members could be expected to have broad transcriptional effects.

**Figure 1:**
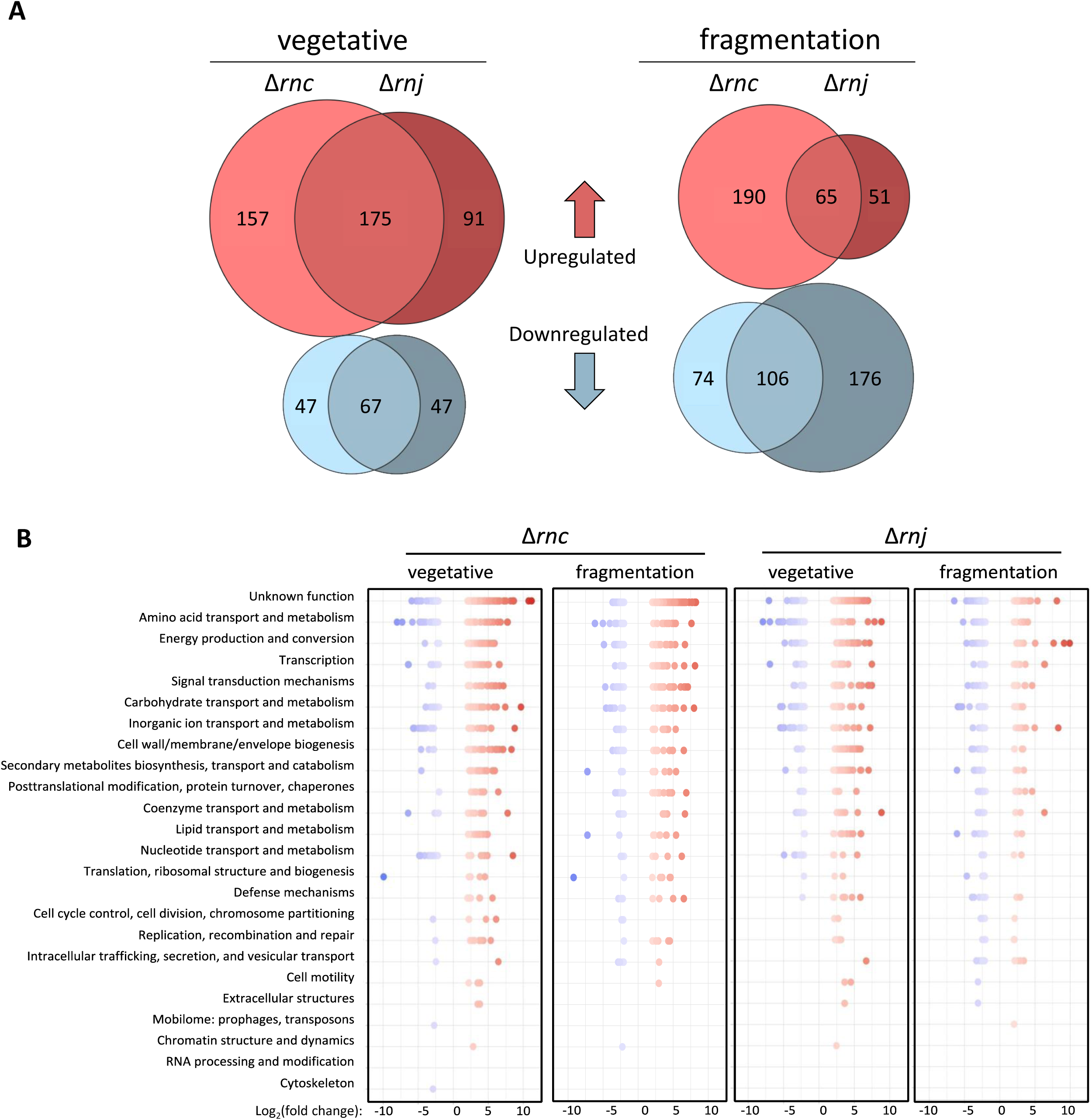
Global shifts in transcript abundance in RNase III and RNase J mutant strains. **(A)** Eulerr plot representing the number of significantly differentially expressed genes shared between RNase III and RNase J mutants, during vegetative and fragmentation stages of development. **(B)** COG analysis for significantly differentially expressed genes, where these genes are sorted into functional categories. Y-axis is ordered by average abundance across all samples.

### Regulators of multicellular development are differentially expressed in RNase mutants

RNases have been previously implicated in the regulation of morphological differentiation in *Streptomyces*. For example, AdpA is a transcriptional regulator that controls the expression of hundreds of genes involved in reproductive growth. In *S. coelicolor* the *adpA* transcript is significantly upregulated in an RNase III mutant and is cleaved by this enzyme *in vitro* [24]. Contrary to what was observed in *S. coelicolor,* we found that *adpA* transcript levels in *S. venezuelae* were not increased in the Δ*rnc* background. Instead, they were less abundant in the Δ*rnc* mutant compared with the wild type during vegetative growth and hyphal fragmentation (although these differences were not statistically significant), suggesting that *adpA* is unlikely to be a direct target of RNase III in *S. venezuelae.* This suggests that RNase III may be co-opted for species-specific functions depending on the abundance and availability of its target transcripts, and their respective structures *in vivo*.

While we observed no significant change in *adpA* transcript levels, additional analyses revealed that many other developmental genes were significantly differentially expressed in both the Δ*rnc* and Δ*rnj* mutants. For example, we found that *bldM* and *bldN* were highly upregulated during vegetative growth with respect to the wild type. These are both known targets of the master regulator BldD [25, 26], and indeed, most members of the BldD regulon were upregulated during vegetative growth in both mutant strains, although the expression of *bldD* itself was not significantly altered (**Figure 2A**). BldD activity is controlled by cyclic-di-GMP (c-di-GMP), where this molecule promotes BldD dimerization, enabling DNA binding and target gene repression [27–29]. Since we observed changes in the BldD regulon but not *bldD* levels, we therefore asked whether any diguanylate cyclase-(responsible for c-di-GMP synthesis) or phosphodiesterase-(responsible for c-di-GMP degradation) encoding genes were differentially expressed in either the Δ*rnc* or Δ*rnj* mutant strains. We found that many of these genes had altered transcript abundances (**Figure 2B**). During vegetative growth, the diguanylate cyclase-encoding genes *cdgE* and *cdgC* were upregulated in the Δ*rnc* and Δ*rnj* mutants, respectively. During hyphal fragmentation, the diguanylate cyclase-encoding gene *cdgC* and the phosphodiesterase-encoding gene *rmdB* were downregulated in both mutant strains, whereas *cdgF* and *vnz_36075* (encoding a bifunctional diguanylate cyclase/phosphodiesterase and putative diguanylate cyclase, respectively) were upregulated in both mutant strains. *cdgE* and *cdgD* were also upregulated during hyphal fragmentation in the Δ*rnc* and Δ*rnj* mutant, respectively [30]. This collectively suggested that c-di-GMP pools may be altered in the RNase mutant strains. The observed upregulation of BldD regulon members suggests that this repressor is less active, implying that there may be reduced concentrations of c-di-GMP available (either globally or locally) to promote its dimerization and repressive function.

**Figure 2:**
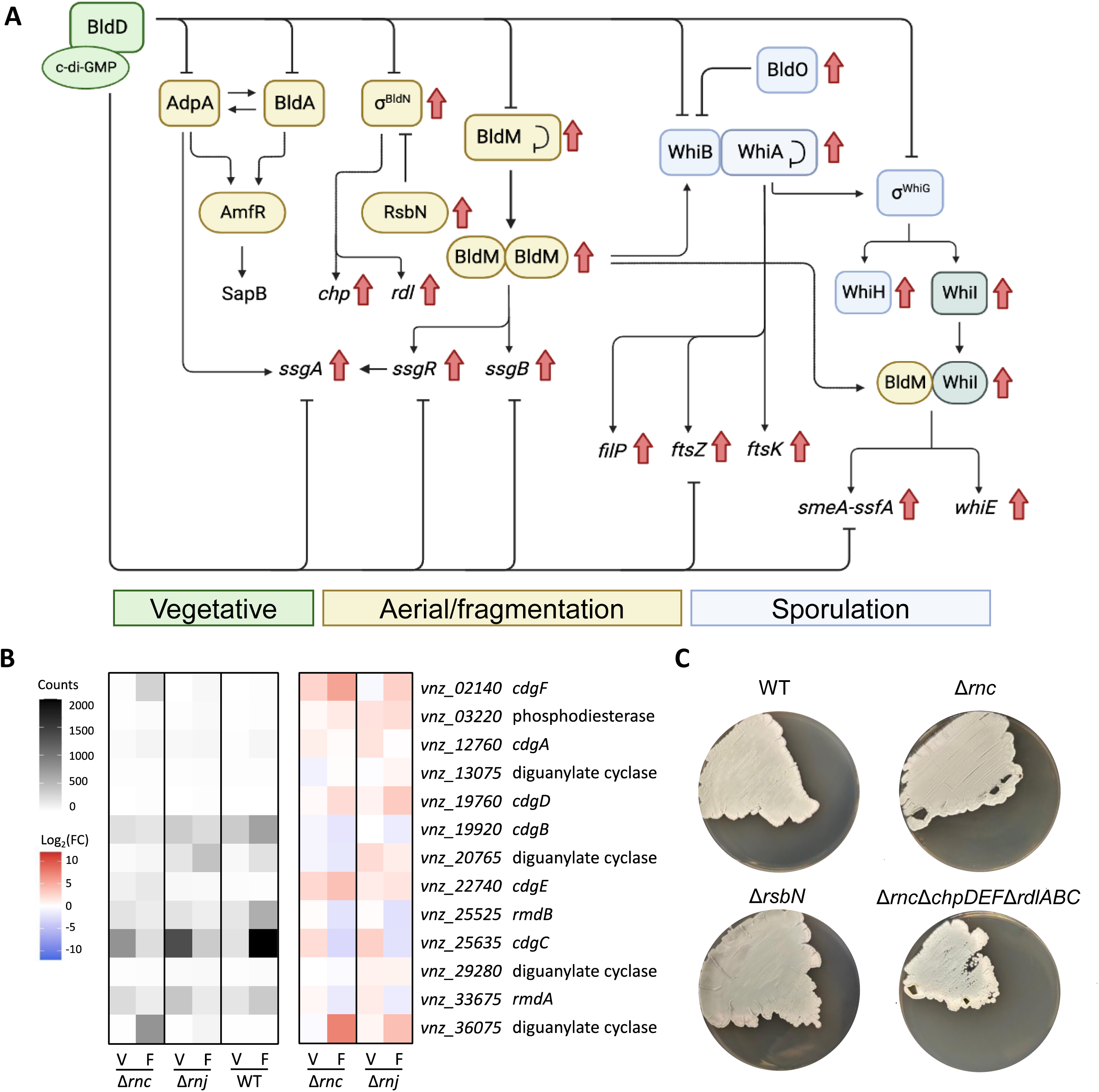
Regulators of multicellular development are differentially expressed in RNase mutants. **(A)** Schematic of the BldD regulon, colored by stage of development where the associated gene product is typically active. Red arrows indicate genes that are significantly upregulated in both the RNase III and RNase J mutant strains, relative to the wild type. **(B)** Differential expression of diguanylate cyclases and phosphodiesterases in *S. venezuelae.* Normalized counts for each strain (normalized across biological replicates) are presented on the left, and Log_2_(fold change) for these counts between the wild type and either RNase mutant is presented on the right. V = vegetative, F = fragmentation. **(C)** Phenotypic comparisons between wild type and *rnc* mutant strains (top left and right, respectively), relative to the *rnc* mutant strain overexpressing the chaplin/rodlin genes (*rsbN* mutant; bottom left), and the *rnc* mutant strain lacking several chaplin and rodlin genes (bottom right).

Included amongst the differentially expressed genes within the (indirect) BldD regulon were the rodlin and chaplin genes, which were highly upregulated in both Δ*rnc* and Δ*rnj* mutant strains. The function of the rodlin and chaplin proteins is to coat the surface of the aerial hyphae, where they confer these hyphae with hydrophobic properties that promote their escape of surface tension and enable their extension into the air [31–36]; these genes are under the direct control of sigma BldN (σ^BldN^). The precocious upregulation of these morphogenetic protein-encoding genes during vegetative growth did not lead to premature reproductive growth for either the Δ*rnc* or Δ*rnj* mutants. Instead, we observed that the Δ*rnj* mutant was significantly delayed in sporulation. We did, however, note that the *rnc* mutant had a tendency to ‘peel’ away from the growth substrate on solid medium, and we wondered if this phenotype may be a result of enhanced chaplin/rodlin production, given the surfactant capabilities of both these protein family members. To investigate this possibility, we generated two *S. venezuelae* strains: one where the rodlins and chaplins were expected to be overexpressed, and another where most of the chaplins and rodlins were deleted in the Δ*rnc* mutant background. To achieve overexpression, we generated an Δ*rsbN* mutant, where *rsbN* encodes the σ^BldN^-specific anti-sigma factor. *rsbN* deletion leads to hyperactivity of σ^BldN^, which in turn increases the expression of the rodlin and chaplin genes [37]. In parallel, we generated a Δ*rnc*Δ*chpDFG*Δ*rdlAB* mutant strain, where many of the rodlin (two of three) and chaplin (three of seven) genes were deleted in the Δ*rnc* mutant background. We had predicted that if the rodlins and chaplins were responsible for the *rnc* peeling phenotype, then the ‘peeling’ aspect of the Δ*rnc* mutant phenotype would be recapitulated in the Δ*rsbN* strain and resolved in the Δ*rnc*Δ*chpDFG*Δ*rdlAB* strain. Instead, we found that deleting the rodlin and chaplin genes did not alter the peeling characteristics of the Δ*rnc* mutant phenotype (**Figure 2C**), nor was an equivalent peeling phenotype seen for the Δ*rsbN* strain, suggesting that this phenotype was not a result of precocious rodlin and chaplin production.

### Regulators of nutrient acquisition are differentially expressed in RNase mutants

Given that the initiation of *Streptomyces* reproductive growth is thought to occur in response to nutrient starvation, we considered whether the RNase mutants may be dysregulated in their nutrient uptake or metabolism. In looking at different nutrient acquisition pathways, we found that many genes involved in nitrogen assimilation and phosphate uptake were differentially expressed in both the Δ*rnc* and Δ*rnj* mutant strains relative to wild type.

Genes related to nitrogen assimilation were upregulated during reproductive growth, particularly in the Δ*rnj* mutant; there was little expression seen during vegetative growth for any strain. Notably, nearly all of these differentially affected genes were part of the GlnR regulon (**Figure 3A**). Under nitrogen-limited growth conditions, GlnR activates the expression of genes involved in nitrogen assimilation, including those encoding nitrate and ammonium transporters, assimilatory nitrate/nitrite reductases, and glutamine synthetases [38, 39]. Comparing transcript abundance within each mutant over time showed that these higher levels of gene expression continued into the sporulation phase for both Δ*rnc* and Δ*rnj* mutants (**Figure 3A**). These increased transcript levels were unlikely to be the result of increased *glnR* expression, as its transcript levels did not change significantly in either mutant relative to the wild type, at any time point (**Figure 3A**). In contrast to the nitrogen-associated regulon, the genes encoding the PhoRP two-component system were downregulated during vegetative growth (**Figure 3B**). This system responds to phosphate limitation by increasing expression of genes involved in phosphate uptake [40–43]. Under low phosphate growth conditions, the PhoR kinase phosphorylates and activates the PhoP response regulator. We observed that several PhoP target genes were expressed at lower levels in the RNase mutants compared with the wild type during vegetative growth (and fragmentation for the *rnj* mutant), including the *pst* operon, which encodes a phosphate ABC transporter; *phoD*, which encodes a secreted alkaline phosphatase; and *phoU*, which encodes a phosphate transport system regulatory protein (**Figure 3B**).

**Figure 3:**
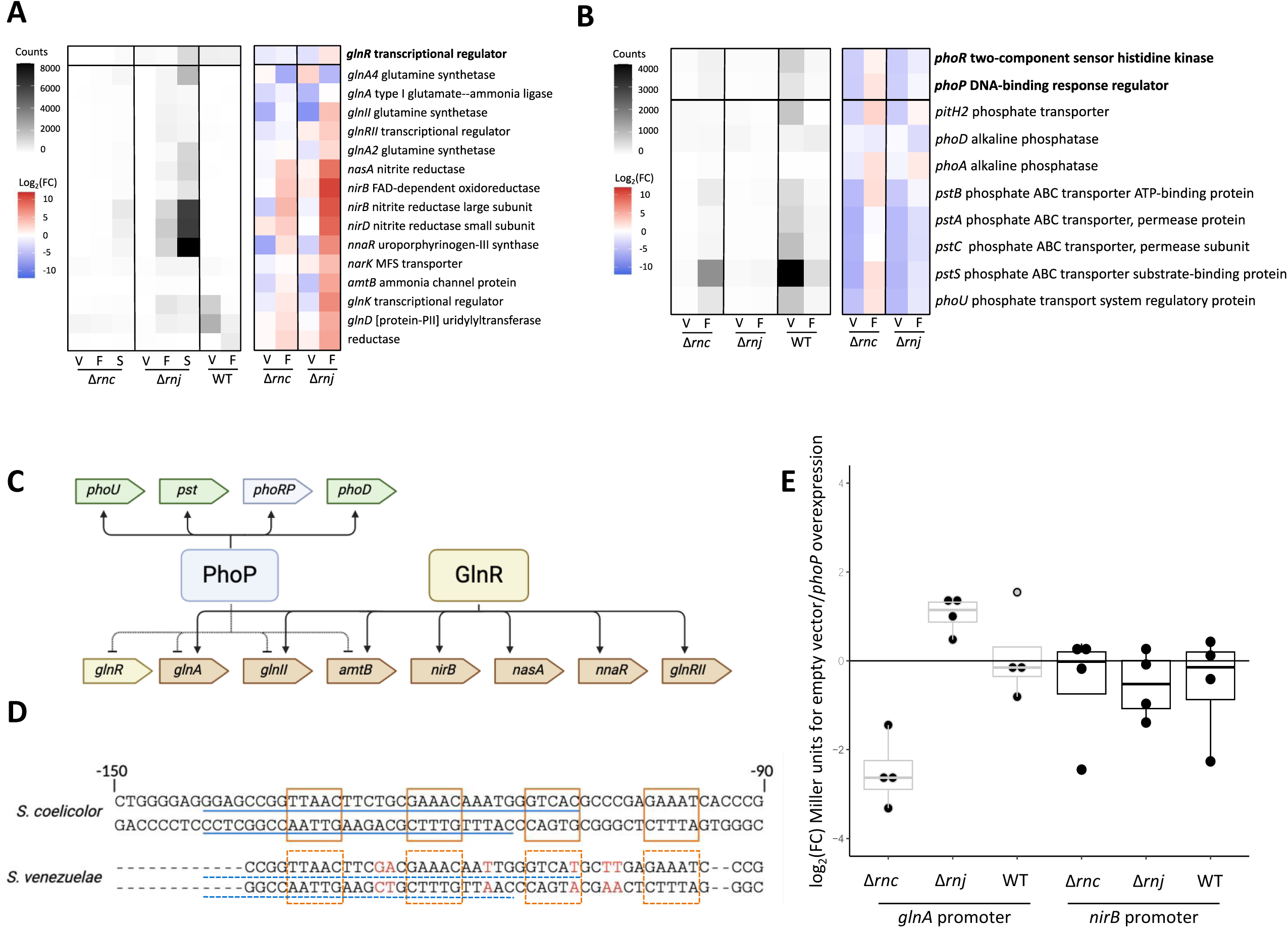
Dysregulation of nitrogen assimilation and phosphate uptake-related genes in RNase mutants. **(A)** Normalized counts (averaged across biological replicates) for GlnR regulon members are presented for RNase mutants at three stages of development, and for wild type at two stages of development (left). Log_2_(fold change) for these counts between the wild type and either RNase mutant is presented on right. V = vegetative, F = fragmentation, S = sporulation. **(B)** Left: normalized counts for the PhoRP two-component system-encoding genes and their associated regulon for each strain (normalized across biological replicates). Right: Log_2_(fold change) for these counts between the wild type and either RNase mutant. **(C)** Schematic illustrating the known crosstalk between GlnR and PhoP. **(D)** Overlapping GlnR binding sites (orange boxes) and PhoP binding sites (blue line) for the *glnA* promoter region in *S. coelicolor* and their conservation in *S. venezuelae* (putative binding sites in dashed boxes/lines). Nonconserved resides are coloured in red. **(E)** Fold change was calculated for each biological replicate, where the difference between promoter activity in a strain carrying an empty vector and a strain carrying the *phoP* phosphomimetic construct was calculated and log_2_ transformed, for the each of two promoters (*glnA* and *nirB*), in wild type, *rnc* and *rnj* mutant strain backgrounds.

The phosphate uptake and the nitrogen assimilation regulatory networks are interconnected, with PhoP competing with GlnR for binding to the promoters of several nitrogen assimilation genes (**Figure 3C**) [42, 44, 45]. The regulatory effects of GlnR and PhoP on the nitrogen assimilation genes have also been reported to be inversely correlated, with GlnR activating their expression and PhoP repressing them. In *S. coelicolor,* shared GlnR/PhoP target promoters include *glnR*, *glnA* and *glnII* (glutamine synthetases), as well as *amtB* (ammonium transporter) [45]. If PhoP was less abundant in the *S. venezuelae* RNase mutants, as suggested by the RNA-sequencing data, then GlnR may have less competition in binding their shared targets, leading to a concomitant increase in transcription of these shared target genes.

We therefore sought to investigate the contribution of *phoRP* downregulation to the Δ*rnc* and Δ*rnj* mutant transcriptomes. We reasoned that if reduced competition between GlnR and PhoP was responsible for the upregulation of downstream genes (*e.g., glnII*, *glnA*, and *amtB*), then overexpressing *phoP* should return transcript levels of these genes in the mutants to near wild type levels. To test this, we generated a *phoP* phosphomimetic overexpression construct [43, 46] and introduced it into Δ*rnc*, Δ*rnj*, and wild type *S. venezuelae* strains. To monitor the expression of nitrogen-assimilation-related genes, we employed a GusA reporter system and introduced the reporter constructs into our *phoP* overexpression and control strains. We monitored the activity of the *nirB* promoter as a negative control (not reported to be co-regulated by GlnR and PhoP) and the *glnA* promoter as our candidate regulon member, as it is targeted by both PhoP and GlnR in *S. coelicolor* [45] and has conserved binding sites in *S. venezuelae* (**Figure 3D**). We measured the promoter activity of *nirB* and *glnA* in strains with an empty plasmid control and compared this to the same strain with the *phoP* phosphomimetic overexpression plasmid. As expected, *nirB* reporter activity was unaffected by *phoP* overexpression (**Figure 3E**). Conversely, *glnA* promoter activity was reproducibly reduced in the Δ*rnc* mutant carrying the *phoP* phosphomimetic overexpression construct compared with its control strain (empty plasmid-carrying Δ*rnc* mutant) (**Figure 3E**). The *phoP* overexpression construct had no effect, however, on *glnA* promoter activity in the Δ*rnj* mutant or wild type strains. This suggested that the Δ*rnc* mutant may be more sensitive to changes in PhoP-mediated regulation than either the wild type or the Δ*rnj* mutant.

### PhoP contributes to precocious chloramphenicol production in Δrnc but not Δrnj

Beyond the effect of PhoP on nitrogen metabolism, there can also be significant crosstalk between PhoP and specialized metabolic regulators. This has been seen in *S. coelicolor,* where PhoP and AfsR compete for binding to the *afsS* promoter, and AfsS in turn impacts the expression of the actinorhodin and undecylprodigiosin biosynthetic clusters [47]. Given the disruptions we found in the PhoRP regulon in our RNase mutants, we sought to assess whether specialized metabolic gene expression in the Δ*rnc* and Δ*rnj* mutant strains differed from that of the wild type.

In both RNase mutant strains, we identified genes having altered transcript abundance in five biosynthetic gene clusters. These clusters were predicted to direct the production of chloramphenicol, melanin, an uncharacterized lanthipeptide, an unknown non-ribosomal peptide, and a terpene (based on antiSMASH detection [48]). For the affected genes in each cluster, precocious expression was seen in the two RNase mutant strains, with transcripts levels being significantly higher than in the wild type during vegetative growth. We set out to test if the increased expression of biosynthetic gene clusters also resulted in increased cluster product levels, and if so, whether this increased specialized metabolite production stemmed from *phoRP* downregulation. For this, we focused on the chloramphenicol biosynthetic cluster due to the strong upregulation of its genes in both mutants (**Figure 4A**) and its well-characterized biosynthetic pathway [49–52]. We quantified chloramphenicol abundance during vegetative growth in the wild type, Δ*rnc* mutant, and Δ*rnj* mutant strains of *S. venezuelae*, with each carrying either our *phoP* phosphomimetic overexpression construct or an empty vector as a control. We found that both Δ*rnc* and Δ*rnj* mutants produced significantly higher levels of chloramphenicol compared with the wild type, and that chloramphenicol levels were higher in the Δ*rnj* mutant than the Δ*rnc* mutant (**Figure 4B**). We further observed that in the Δ*rnc* mutant strain, *phoP* overexpression reduced chloramphenicol production to roughly wild type levels, indicating that *phoRP* downregulation likely contributed to the increased antibiotic production observed in the Δ*rnc* mutant strain. Our findings here mirrored our observation that PhoP affected *glnA* promoter activity in the Δ*rnc* mutant but not the Δ*rnj* mutant. Indeed, *phoP* overexpression did not significantly impact chloramphenicol levels in the Δ*rnj* mutant relative to its control strain (**Figure 4B**). While there was a trend observed for less chloramphenicol production in the Δ*rnj* mutant following *phoP* overexpression, this reduction did not approach wild type levels; therefore, we could not attribute the increased antibiotic production in the Δ*rnj* mutant solely to *phoP* downregulation. It seems that while the Δ*rnc* and Δ*rnj* mutants have similar transcriptional and metabolic profiles, different mechanisms appear to drive the observed effects.

**Figure 4:**
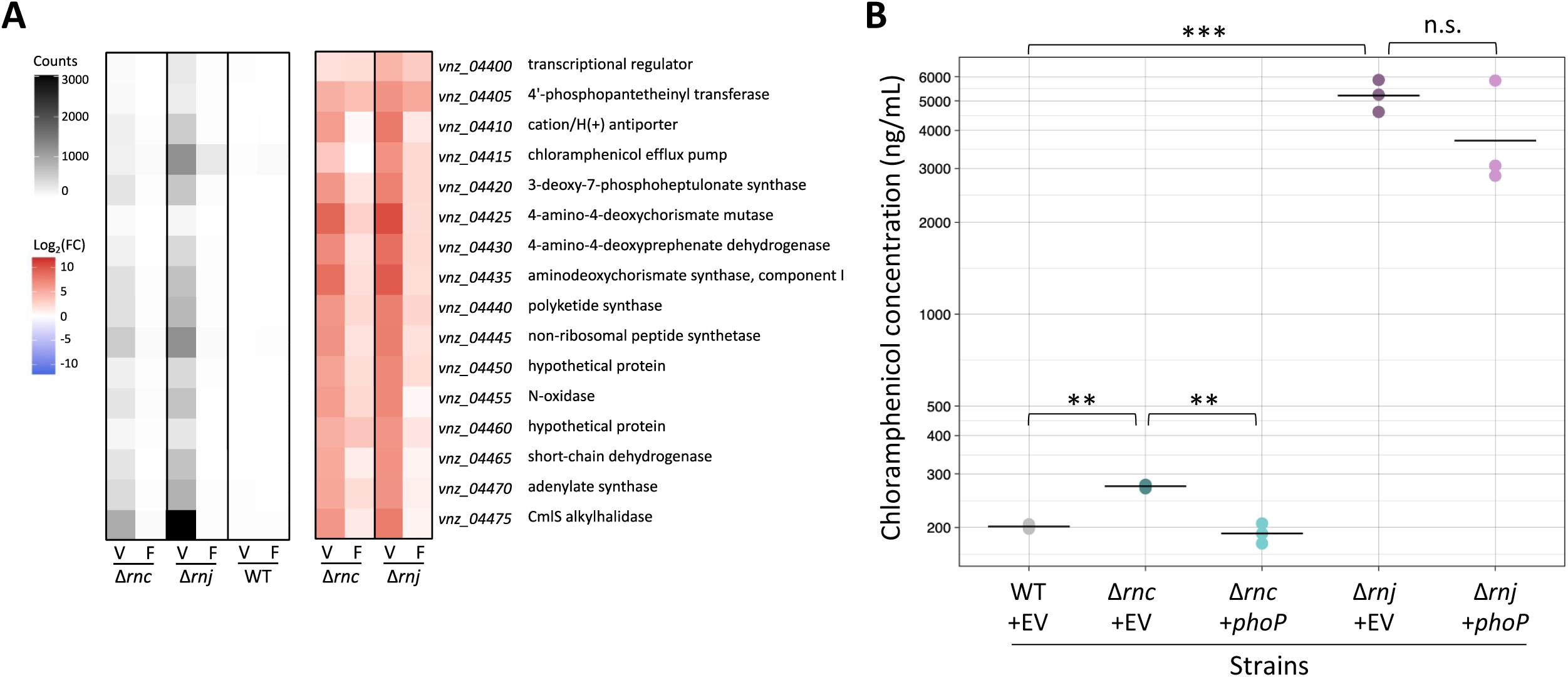
Upregulation of the chloramphenicol biosynthetic cluster is associated with precocious chloramphenicol production in both RNase mutant strains. **(A)** Chloramphenicol biosynthetic gene expression was assessed in Δ*rnc* and Δ*rnj* mutants relative to the wild type during vegetative growth. Normalized counts for each strain (normalized across biological replicates) are presented on the left, and the Log_2_(fold change) for these counts between the wild type and either RNase mutant is presented on the right. V = vegetative, F = fragmentation. **(B)** Quantification of chloramphenicol production levels in wild type (carrying an empty vector (EV) as a control), compared with RNase mutants carrying either an empty vector (EV) or vector with an overexpression *phoP* phosphomimetic variant (+*phoP*).

### phoU is directly targeted by RNase III

Having established that multiple aspects of the Δ*rnc* mutant phenotype were due to reduced expression of the *phoRP* genes, we next wanted to better understand the connection between RNase III and this operon. In *E. coli* the *phoP* transcript can be bound by a small, *cis-* encoded antisense RNA, with the resulting double-stranded complex being targeted for degradation by RNase III [53]. In *S. venezuelae,* we noted low level antisense reads mapping to the *phoRP* operon (**Figure 5A**; red arrow), and so we wondered if *phoP* may be targeted by RNase III in *S. venezuleae* in a similar way as in *E. coli*.

**Figure 5:**
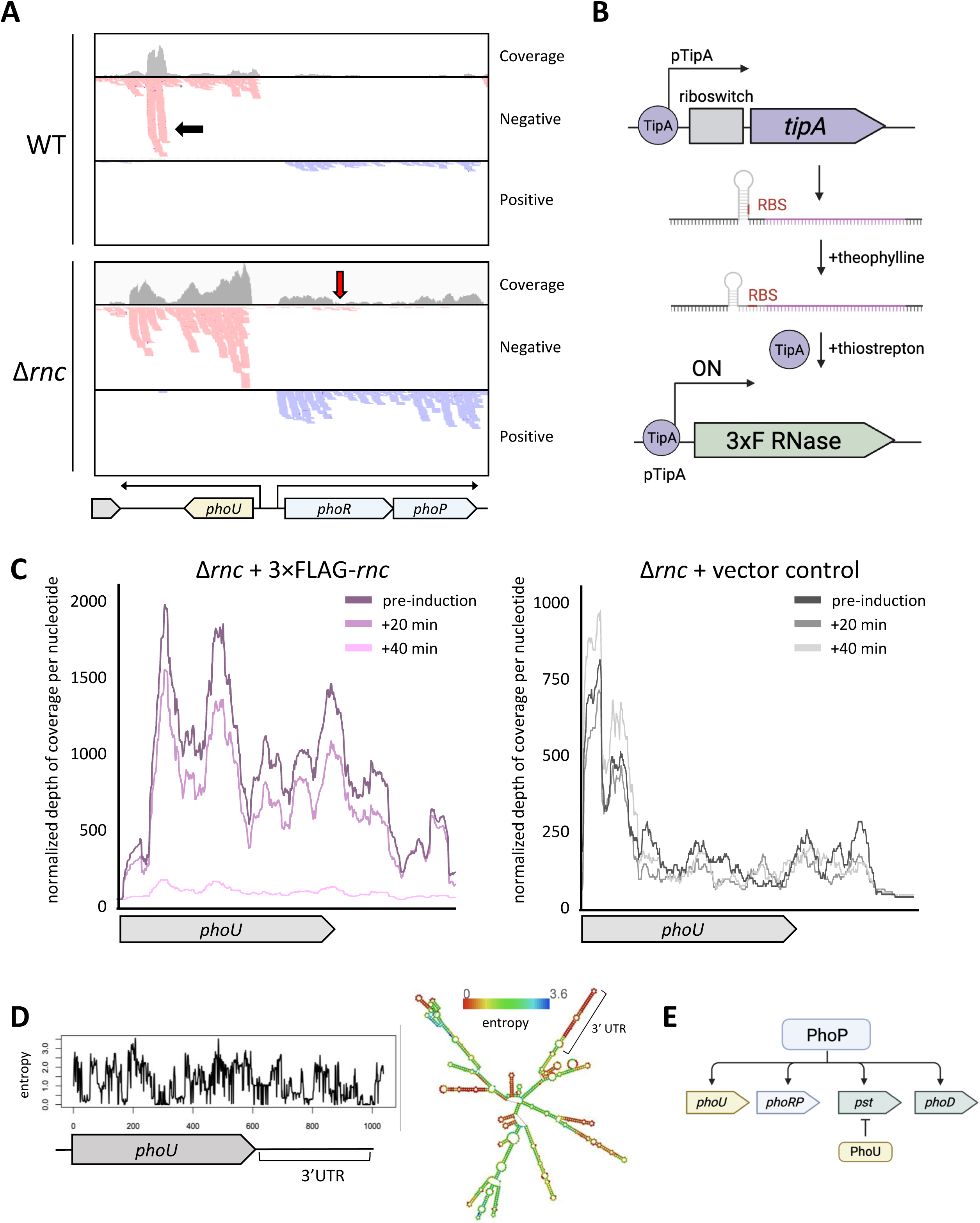
PhoU transcript is directly targeted by RNase III. **(A)** Reads aligning to the *phoRP* and *phoU* region from the wild type and RNase III mutant samples (during fragmentation), including a pileup of reads aligning to the 3’ UTR of *phoU* (black arrow), and a low level of antisense reads aligning to the *phoR* gene in the RNase III mutant background (red arrow). **(B)** Schematic for the transcriptionally- and translationally-acting inducible system. Addition of theophylline relieves translational repression, where the ribosome binding site (RBS) upstream of *tipA* becomes accessible. Subsequent addition of thiostrepton increases the affinity of TipA for its own promoter sequence, which is upstream of *rnc* fused to a 3×FLAG tag N-terminal sequence. **(C)** Reads aligning to *phoU,* before and after RNase III-induction. Normalized depth of coverage per nucleotide is plotted over the *phoU* coding region and into the 3’UTR in the experimental sample (carrying the 3 × FLAG tagged RNase III; left) and the control sample (lacking the inducible RNase III construct; right). Depth of coverage/nucleotide was normalized by library size, and a loess regression trendline is presented (span = 0.1 was used to account for high /nucleotide variability; representative biological replicate is presented here). The scripts used for normalization and generating coverage plots are provided in supplementary materials. **(D)** *phoU* transcript has highly structured 3’UTR based on RNAfold predictions (right), and this region is associated with lower entropy compared with the coding sequence (left). **(E)** Schematic indicating a role for PhoU in a negative feedback loop within the PhoP regulon, where it represses the expression of several PhoP-activated genes, including the *pst* operon.

To identify transcripts that were directly targeted for degradation by RNase III, we designed an inducible system that allowed us to control *rnc* expression, reasoning that direct targets would be degraded more rapidly than indirect targets. To minimize leaky expression from our uninduced construct, we employed a two-tiered inducible system, encompassing a translationally-acting theophylline-responsive riboswitch controlling the *tipA* gene, and a thiostrepton-inducible promoter that could be induced by thiostrepton-bound TipA (**Figure 5B**). We then cloned a FLAG-tagged *rnc* gene downstream of the thiostrepton-inducible promoter, introduced the resulting construct into the *rnc* mutant strain, and confirmed robust 3×FLAG-RNase III production after adding the theophylline and thiostrepton inducers (**Supplementary Figure 1**). We collected cells prior to induction, and at 20- and 40-minutes post-induction (when we observed strong induction of RNase III; **Supplementary Figure 1**), and isolated RNA from these samples, alongside a Δ*rnc* strain bearing an empty inducible vector to control for the effects of theophylline and thiostrepton addition. These samples were then subjected to RNA-sequencing. We focused on genes that were both significantly downregulated in the induced samples relative to the uninduced, and whose transcript levels were unaffected by inducer addition in the control strain. Of the RNase III targets we identified, *phoU* was of particular interest. Its transcript levels decreased dramatically post-RNase III induction and were generally unchanged in the control when comparing uninduced and induced samples (**Figure 5C**). In *S. venezuelae, phoU* is expressed divergently from the *phoRP* operon (**Figure 5A**; bottom). We observed that its transcription continued beyond the end of the *phoU* coding sequence, yielding a long 3ʹ UTR. A segment of this UTR was more abundant in the wild type compared with the Δ*rnc* mutant, suggesting that this may be a stable product that accumulates following RNase III cleavage (**Figure 5A**; black arrow). Whether this product has any function in the cell remains to be determined. In analysing the structure of the *phoU* transcript using the RNAfold program [54], we found the 3ʹ UTR was predicted to be highly structured (**Figure 5D**), suggesting that the many predicted stem-loops could serve as substrates for direct RNase III-mediated cleavage.

### Ribosomal protein transcripts are directly targeted by RNase J

While the PhoP-mediated effects appeared to be specific to RNase III and not RNase J, we remained intrigued by the congruence of the transcription profiles for both Δ*rnc* and Δ*rnj* mutant strains. We wondered what transcripts were directly targeted by RNase J and whether these could provide insight into the affected pathways that differ when compared with the Δ*rnc* mutant (*e.g.,* chloramphenicol biosynthesis being upregulated in both mutants, but via apparently different pathways). We generated an RNase J-inducible construct using the theophylline/thiostrepton-inducible system described above and introduced this (alongside an empty plasmid control) into the Δ*rnj* mutant in *S. venezuelae.* We first confirmed that the 3×FLAG-RNase J was both functional and could be effectively induced (**Supplementary Figure 1B**). We then conducted an equivalent induction experiment as described above for RNase III. In focusing on transcripts that decreased post-induction in the RNase J experimental samples but not in the control, we found no target transcripts that could easily explain the Δ*rnj* mutant phenotypes described so far. Considering the list of RNase J target transcripts, it was instead immediately striking that half of them (14/28) encoded ribosomal proteins (**Table 1**). These transcripts all decreased dramatically (between 3- and 30-fold) post-RNase J induction compared to the control samples, where no change in abundance was detected (**Table 1**; **Figure 6**). Notably, this effect was not seen following RNase III induction, indicating that it is specific to RNase J. Ribosomal proteins are generally understood to autogenously regulate their own mRNA transcripts by binding to structured regions in the associated 5′ UTRs [55, 56]. Our observations here suggest a potentially novel mechanism for ribosomal protein control, whereby RNase J exerts post-transcriptional regulation of these transcripts.

**Figure 6:**
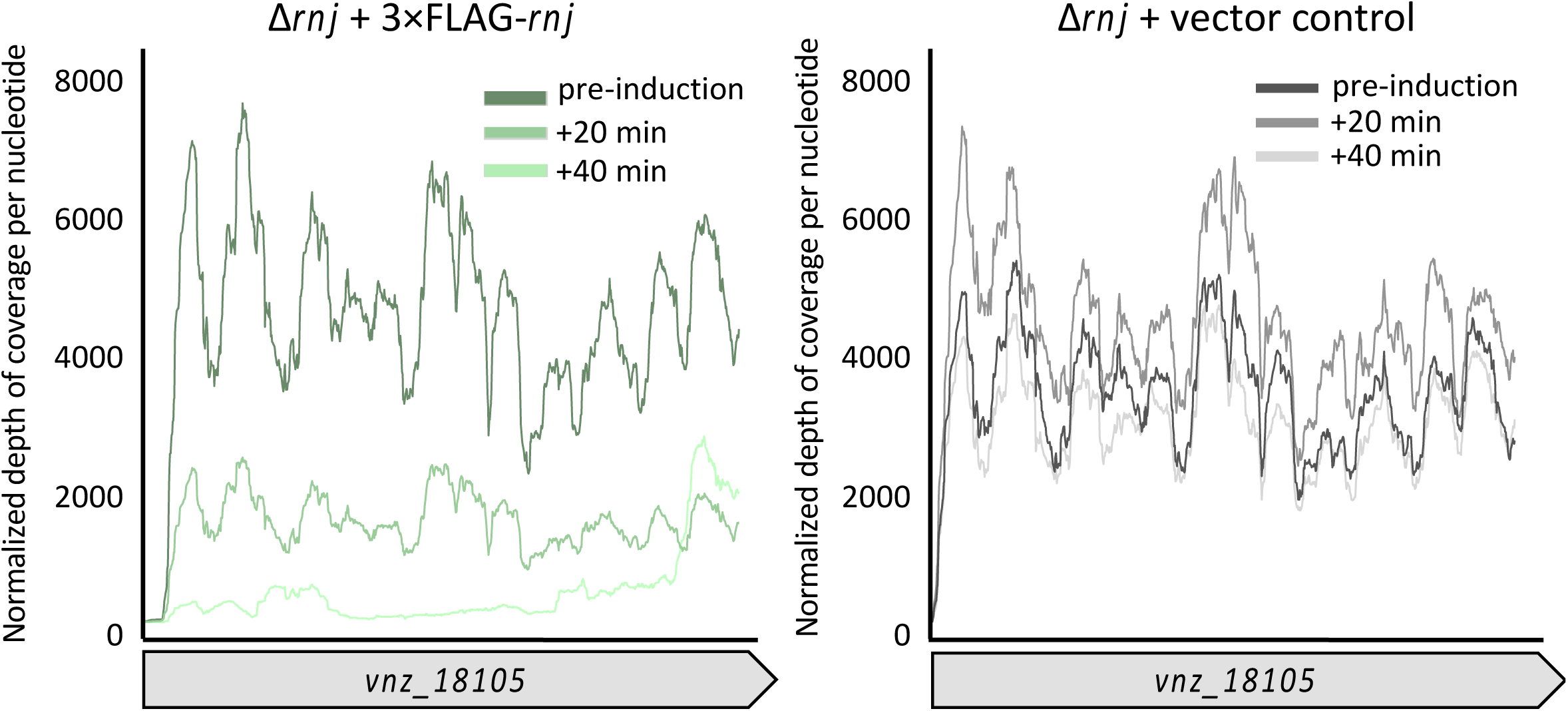
RNase J is involved in regulating translational machinery. *vnz_18105* (encoding a 30S ribosomal protein) is a likely RNase J target gene. Normalized depth of coverage per nucleotide is plotted over the *vnz_18105* coding region in the experimental sample (carrying the 3×FLAG tagged RNase J; left) and the control sample (lacking the inducible RNase J construct; right). Depth of coverage/nucleotide was normalized by library size, and a loess regression trendline is presented (span = 0.1 was used to account for high /nucleotide variability; representative biological replicate is presented here).

**TABLE 1.** Genes whose transcripts are directly targeted by RNase J post-induction.

| Gene name and associated product | Fold change pre- to 40-min |
| --- | --- |
| **Ribosomal protein-encoding genes** |  |
| 30S ribosomal protein S4 (*vnz_05370*) | -9.2 |
| 50S ribosomal protein L20 (*vnz_05835*) | -14.2 |
| 30S ribosomal protein S1 (*vnz_07970*) | -3.8 |
| 30S ribosomal protein S20 (*vnz_11455*) | -9.3 |
| 30S ribosomal protein S6 (*vnz_18105*) | -15.9 |
| 50S ribosomal protein L11 (*vnz_21445*) | -9.3 |
| 30S ribosomal protein S12 (*vnz_21600*) | -14.5 |
| 30S ribosomal protein S17 (*vnz_21685*) | -31.3 |
| 30S ribosomal protein S11 (*vnz_21775*) | -10.1 |
| 50S ribosomal protein L13 (*vnz_21805*) | -16.5 |
| 50S ribosomal protein L32 (*vnz_26035*) | -3.8 |
| 50S ribosomal protein L19 (*vnz_26165*) | -9.3 |
| 30S ribosomal protein S2 (*vnz_26230*) | -4.9 |
| 30S ribosomal protein S15 (*rpsO*; *vnz_26635*) | -9.6 |
| **Other targets** |  |
| elongation factor P (*vnz_05295*) | -4.5 |
| peptidylprolyl isomerase (*vnz_06015*) | -3.0 |
| hypothetical protein (*vnz_07005*) | -10.4 |
| leucine tRNA ligase (*vnz_11510*) | -2.3 |
| hypothetical protein (*vnz_15735*) | -16.0 |
| aspartate tRNA ligase (*vnz_17530*) | -2.8 |
| translation initiation factor IF-1 (*vnz_21760*) | -6.1 |
| hypothetical protein (*vnz_24805*) | -3.6 |
| acetolactate synthase large subunit (*vnz_25640*) | -2.1 |
| hypothetical protein (*vnz_26030*) | -7.9 |
| RNase III (*rnc*; *vnz_26040*) | -3.3 |
| translation elongation factor Ts (*vnz_26235*) | -4.6 |
| hypothetical protein (*vnz_26610*) | -3.2 |
| PNPase (*pnp*; *vnz_26440*) | -2.4 |

### Co-regulation of specific transcripts by RNase III and RNase J

Reflecting on the role that RNase J may have in regulating translational machinery, we noted that this was another pathway where there was congruence between RNase III and RNase J, having previously shown that RNase III was involved in 16S rRNA maturation [22]. Beyond the effects of these ribonucleases on discrete ribosome components, we also observed that *ssrA* [a transfer-messenger RNA (tmRNA)] was upregulated in the Δ*rnc* mutant, although not in the Δ*rnj* mutant (**Supplementary Tables S1-S4**). tmRNA molecules function to rescue stalled ribosomes [57], and upregulation of tmRNA in the Δ*rnc* mutant suggested there may be increased numbers of stalled ribosomes in the absence of RNase III. The shared transcriptional profiles observed for our Δ*rnc* and Δ*rnj* mutant strains, which were markedly distinct from that of the wild type strain, may therefore stem from (independent) defects in translation, given the direct coupling of transcription and translation in many bacteria [58–64].

An alternative/additional explanation for the similar transcriptomic shifts observed in the Δ*rnc* and Δ*rnj* mutant strains could be that some transcripts are directly targeted by both RNase III and RNase J. Mechanistically, there is the potential for these ribonucleases to cooperatively degrade target transcripts: RNase J preferentially recognizes and degrades transcripts having a 5ʹ monophosphate, which can arise through RNase III cleavage (**Figure 7A**). There is at least one previously reported instance of their coordinated activity in *Staphylococcus aureus*, where these ribonucleases act to cooperatively cleave the methionine biosynthesis T-box riboswitch [65]. Given this precedent, we sought to investigate whether any target transcripts were direct targets for degradation by both RNase III and RNase J in *S. venezuleae*.

**Figure 7:**
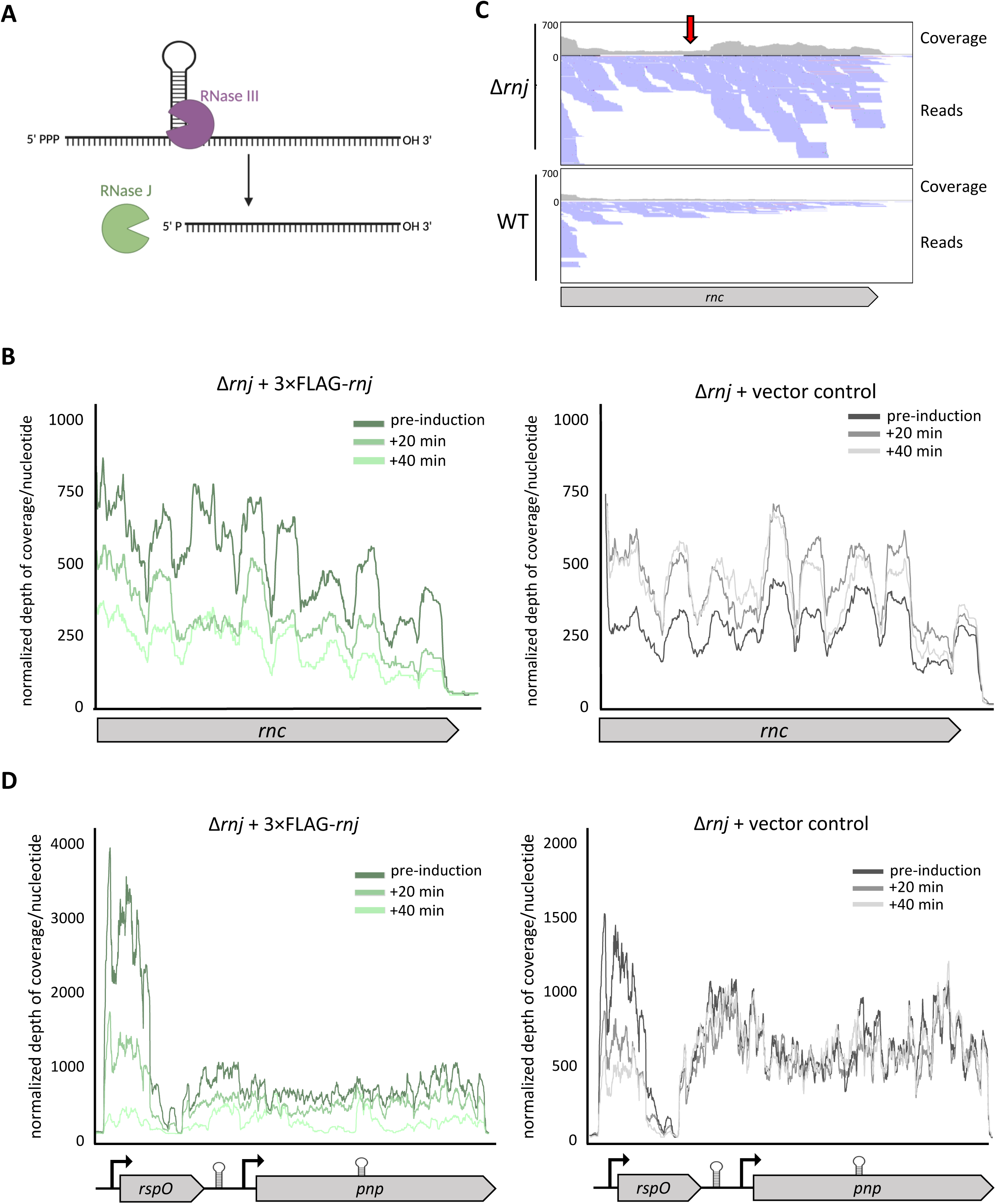
RNase III and RNase J may cooperatively degrade specific transcripts. **(A)** Schematic for mechanism of RNase III and RNase J cooperative degradation. Initial endoribonucleolytic cleavage by RNase III yields a 5ʹ monophosphorylated transcript, which is preferentially recognized by RNase J for exoribonucleolytic degradation. **(B)** Reads aligning to *rnc* are presented as normalized depth of coverage per nucleotide plotted over the *rnc* coding region in the experimental sample (carrying the 3×FLAG tagged RNase J; left) and the control sample (lacking the inducible RNase J construct; right). Depth of coverage/nucleotide was normalized by library size, and a loess regression trendline is presented (span = 0.1 was used to account for high /nucleotide variability; representative biological replicate is presented here). The scripts used for normalization and generating coverage plots are provided in the supplementary materials. **(C)** Reads aligning to the *rnc* coding region from the wild type and RNase J mutant samples (during fragmentation). The red arrow indicates a predicted internal cleavage site that could then serve as a substrate for RNase J (given the read pileup aligning downstream of this site in the RNase J mutant but not wild type). **(D)** Reads are aligned to *rspO* and *pnp,* with normalized depth of coverage per nucleotide plotted over the *rspO-pnp* coding region in the experimental sample (carrying the 3×FLAG tagged RNase J; left) and the control sample (lacking the inducible RNase J construct; right).

Our RNase III- and RNase J-induction experiments did not identify any shared target transcripts. We did, however, observe that two previously described RNase III targets [66, 67] appeared to be directly cleaved by RNase J. In many bacteria, including *E. coli* and *S. coelicolor*, RNase III exerts autoregulatory control by cleaving a stem-loop structure within its *rnc* transcript [11, 67]. We found that the *rnc* coding region was rapidly degraded in *S. venezuelae* in our RNase J-induced samples (**Figure 7B**). Our original RNA sequencing data also revealed that the terminal end of the *rnc* transcript increased in abundance in the Δ*rnj* mutant, suggesting that this enzyme may function to degrade the truncated transcript after internal RNase III-mediated cleavage (**Figure 7C**). It has also been reported previously in *S. coelicolor,* that RNase III can degrade the *rspO-pnp* transcript [11, 66]. This operon encodes a ribosomal protein and a polynucleotide phosphorylase (PNPase) that has both exoribonuclease and RNA 3ʹ-polyribonucleotide polymerase activity. The *rspO-pnp* operon was not identified as an RNase III target in our induced experiment, but it was upregulated in the Δ*rnc* mutant strain to levels that were just below our significance threshold. This region too was identified as a direct target of RNase J (**Figure 7D**), suggesting that this operon may also be under the cooperative control of RNase III and RNase J.

## DISCUSSION

### The effect of RNase III and RNase J on *Streptomyces* development

Our transcriptional analyses of the Δ*rnc* and Δ*rnj* mutant strains revealed that these enzymes had important but distinct roles in regulating morphological development and metabolism. These mutants shared similar gene expression trends when compared with the wild type, particularly concerning the dysregulation of genes whose products were involved in reproductive growth and specialized metabolism. Our work suggested that the master regulator BldD was less active in the Δ*rnc* and Δ*rnj* mutant strains given the strong upregulation of the BldD regulon. While there was differential expression observed for multiple diguanylate cyclases (c-di-GMP ‘makers’) and phosphodiesterases (c-di-GMP ‘breakers’), the redundancy of these enzymes makes it challenging to predict the net effect of these changes on global c-di-GMP pools. There is, however, increasing evidence suggesting that c-di-GMP signaling frequently functions on a more local scale [68], and thus the change in expression seen for the c-di-GMP synthases and phosphodiesterases here may serve to alter the local pools of c-di-GMP relative to wild type, with those impacting BldD appearing to be amongst the most profoundly affected.

Our transcriptional data also suggested that unlike in *S. coelicolor, adpA* was not a direct target of RNase III in *S. venezuelae*, at least under the conditions tested here. The precise site of RNase III cleavage has not been mapped for the *adpA* transcript in *S. coelicolor*, but *in vitro* experiments in which *adpA* transcripts were incubated with purified RNase III yielded multiple products [24], suggesting that there may be several RNase III cleavage sites within the *adpA* transcript. RNase activity can be impacted by both the sequence and structure of its target transcripts, and by target accessibility within the cell (*e.g.,* protection by translating ribosomes). The difference in *adpA* targeting between these two *Streptomyces* species suggests that either the *S. venezuelae* transcript adopts an alternative structure compared with its *S. coelicolor* counterpart, and/or is subject to different translational or accessibility controls.

### Connections between RNase III and phosphate uptake

The *phoRP* operon was among the most significantly differentially expressed genes in the Δ*rnc* and Δ*rnj* mutant strains, relative to wild type. In *Streptomyces* species, there is known crosstalk between the PhoRP and the GlnR regulons [69–72], a phenomenon that was also observed here. In probing this connection, we noted that the Δ*rnc* mutant was more responsive to changes in PhoP levels than the Δ*rnj* mutant, with both chloramphenicol abundance and *glnA* expression being impacted by altered PhoP levels in the RNase III mutant strain. These observations may be explained in part, by the direct targeting of *phoU* transcripts by RNase III.

In *S. coelicolor,* PhoU represses several PhoP-activated genes, including *glpQ1* and *pstS* [73]. Given that *phoU* transcription is activated by PhoP binding to the *phoU-phoRP* intergenic region, this is thought to create a negative feedback loop that can dampen the otherwise strong positive effect of *phoRP* expression (i.e. genes that are initially upregulated by PhoP are then repressed by PhoU; **Figure 5E**) [73]. While *phoRP* and the associated regulon were downregulated in both mutant strains during vegetative growth, we observed that *phoU* was upregulated in the Δ*rnc* mutant relative to wild type during reproductive growth, which would be consistent with *phoU* being a direct target of RNase III (**Figure 3B**). No upregulation was observed in the Δ*rnj* mutant at this same developmental stage, and the connection between *phoRP* downregulation and RNase J activity remains unclear. These data suggest that despite observing similar shifts in transcript abundance following the deletion of either *rnc* or *rnj* in *S. venezuelae*, there seem to be fundamental differences in the regulatory networks that contribute to these shared effects.

Beyond our observations for the *phoRP* operon and the connection between RNase III and *phoU*, we did not always see congruence between our different transcriptomic experiments (*e.g.,* developmental time course versus inducible time course). We had expected that transcripts identified as direct RNase targets would be more abundant in the corresponding RNase mutant strain, where those transcripts would presumably be protected from degradation. This was not, however, consistently observed upon the cross-referencing of these datasets. Our inducible system informed which transcripts decreased in abundance upon exposure to increasing concentrations of RNase III or RNase J; any transcript whose abundance decreased under these conditions represented a putative direct target of RNase III or RNase J. Considering the more global Δ*rnc* and Δ*rnj* mutant RNA sequencing, we are unable to rule out the possibility that some transcripts were both directly and indirectly affected by these RNases; we were also unable to exclude the possibility that they were subject to sequential cleavage by multiple ribonucleases. We would further note that we identified the PNPase-encoding transcript as a potential target of both RNase III and RNase J, where PNPase is a promiscuous ribonuclease that contributes to bulk mRNA decay in *Streptomyces* [74]. Our observations here emphasize the complexity associated with RNA turnover in *S. venezuelae*, and the interplay of the many enzymes that influence transcript stability.

### The role RNase III and RNase J in regulating translational machinery

Our work here and previously [22], have suggested that RNase III and RNase J both contribute to the expression/processing/function of translational machinery, but in different ways. Loss of each RNase has been associated with an accumulation of inactive 100S ribosome dimers [22], with the Δ*rnc* mutant having a further increase in free 30S and 50S subunits (and fewer 70S ribosomes) compared with the wild type and Δ*rnj* mutant strains [22]. For the Δ*rnc* mutant, these ribosome assembly defects may be consistent with increased tmRNA levels. tmRNA functions within the trans-translation pathway to recycle stalled ribosomes by promoting their disassociation, and this activity has the potential to increase the relative levels of 30S and 50S ribosomal subunits [75–78].

While RNase III is known to be involved in the pre-processing of 16S rRNA, our data here indicated that RNase J impacts the turnover of ribosomal protein transcripts. It is well-established in *E. coli* [55] and more recently in the *Streptomyces* relatives *Mycolicibacterium smegmatis* and *Mycobacterium tuberculosis* [56], that many ribosomal protein-encoding genes and operons are subject to autoregulatory feedback inhibition at the transcriptional level. Our observations here may imply the existence of an additional level of ribonucleolytic/post-transcriptional control of ribosomal proteins by RNase J.

### RNase mutant phenotypes have qualities of the stringent response

The effect of mutating *rnc* and *rnj* in *S. venezuelae* has broad phenotypic consequences, including alterations in translational capabilities, nitrogen metabolism, and antibiotic production. When considering these impacted processes, we noted many parallels with the initiation of the stringent response [79, 80]. The stringent response is mediated by accumulation of the second messenger (p)ppGpp, and initiates in response to nitrogen limitation and a corresponding drop in both amino acid levels and charged tRNA pools, ultimately leading to an accumulation of stalled ribosomes [79, 81]. We observed strong upregulation of the GlnR regulon in the Δ*rnc* and Δ*rnj* mutants, which may suggest that these strains are experiencing nitrogen limitation; we further observed upregulation of tmRNA in the Δ*rnc* mutant, where tmRNA contributes to the rescue of stalled ribosomes.

Interestingly, in *S. coelicolor* and other *Streptomyces* species, the accumulation of (p)ppGpp is also necessary for the production of some antibiotics [80, 82–84]. This is reminiscent of our observations here, where our RNase mutants exhibited both significant upregulation of multiple biosynthetic gene clusters and increased antibiotic production. This trend was particularly evident in the RNase J mutant, where the greatest upregulation of nitrogen-assimilation related genes and the greatest increase in chloramphenicol (antibiotic) production were both observed.

### Concluding remarks

We have demonstrated that RNase III and RNase J are integral to the coordinated regulation of primary and specialized metabolism in *S. venezuelae*. These ribonucleases have important roles in ensuring the appropriate timing of key developmental and metabolic processes: loss of either RNase resulted in the dysregulation of genes whose products were involved in development, nitrogen assimilation, phosphate uptake, and specialized metabolism. We identified direct targets of these RNases, provided insight into how their catalytic activity contributed to the regulation of phosphate uptake (and consequent antibiotic production), and revealed differences in their contribution to the production and assembly of ribosomal components. We further revealed the potential for RNase III and RNase J to cooperatively degrade specific transcripts in the case of the *rspO-pnp* operon. This operon is highly conserved, and this mechanism of regulation may be similarly conserved across bacteria encoding both RNase III and RNase J. Our work underscores the complex nature of RNase-based regulation and how these enzymes function to collectively enable the appropriate control of essential cellular processes in bacteria.

## MATERIALS AND METHODS

### Strains, plasmids, media, and culture conditions

Strains, plasmids, and primers used in this study are listed in Supplementary Tables S6–S8, respectively. *S. venezuelae* NRRL B-65442 was grown in liquid MYM (1% malt extract, 0.4% yeast extract, 0.4% maltose) for overnight cultivation and on solid MYM (2% agar) for spore stock generation. All *Streptomyces* cultures were grown at 30°C. *E. coli* DH5α was used for general cloning and plasmid preparation. All *E. coli* strains were grown at 37°C on lysogeny broth (LB) or Difco nutrient agar plates, or in LB or super optimal broth liquid medium, supplemented with antibiotics where appropriate to maintain plasmid selection.

### RNA isolation and sequencing to identify differentially expressed genes and direct RNase targets

RNA was isolated, as described previously [85], from *S. venezuelae* cultures grown in liquid MYM medium. For wild type and RNase mutant transcriptomic analyses, two biological replicates were collected for each strain and stage of growth (with the timing of growth phase harvest optimized for each strain). Early/vegetative samples were collected at 10 h for wild type and the Δ*rnc* strain, and at 12 h for the Δ*rnj* strain. Mid-stage/fragmenting samples were collected at 14 h for wild type and the Δ*rnc* strain, and at 18 h for the Δ*rnj* strain. Late-stage/sporulating samples were collected at 20 h for the Δ*rnc* strain and at 24 h for the Δ*rnj* strain.

Strains carrying our thiostrepton-theophylline inducible system (or the empty plasmid control) were sub-cultured to 0.1 OD_600_ and grown at 30°C for 8 h. Theophylline was then added to a concentration of 4 mM to induce TipA translation, after which cultures were grown for an additional 2 h at 30°C before thiostrepton was added to a final concentration of 50 μg/mL to promote expression of FLAG-tagged RNases. Cells were collected at 0 min (pre-induction), and at 20 and 40 min post-thiostrepton addition. Aliquots were then collected for RNA isolation and western blotting.

RNA-sequencing was conducted by the MOBIX facility at McMaster University using paired-end technology on the Illumina MiSeq v3 platform. The data discussed in this publication have been deposited in NCBI’s Gene Expression Omnibus [86] and are accessible through GEO Series accession numbers GSE285999 and GSE286000.

### Differential expression analysis for RNA sequencing samples

In all instances, reads were aligned to the *S. venezuelae* NRRL B-65442 genome (NCBI accession: NZ_CP018074.1) using the BowTie2 program [87] and the aligned files were sorted using SamTools [88]. The number of reads aligning to genomic features were counted using the HTseq-count program [89] for each replicate. The HTseq-count tables were then normalized and used for differential transcript level analysis with the program DESeq2 [90].

For transcriptomic analysis of the RNase mutants compared to wild type *S. venezuelae,* these data were filtered by significance (adjusted p-value < 0.05), expression (BaseMean > 50), and differential expression (log_2_ fold change > |2|). For our inducible experiments we filtered these data by significance (adjusted p-value < 0.05), expression (BaseMean > 50), and differential expression (fold change < −2). We also used these criteria to exclude any genes whose transcript abundance was significantly altered in the plasmid controls between the 0 min and the 40 min samples, and excluded any genes whose transcript abundance increased post-induction.

### Western blotting to detect FLAG-tagged RNases

We used western blotting to confirm the expression of FLAG-tagged proteins in *S. venezuelae* following induction of their expression. Between 40 - 80 μg of total protein from each sample were run on a 12% SDS-denaturing polyacrylamide gel for 1 h at 150 V. The proteins were transferred to a methanol-activated PVDF membrane (Amersham Bioscience) using the Bio-Rad Trans-Blot® Turbo™ Transfer System with 1× transfer buffer (48 mM Tris, 39 mM glycine, 1.28 mM SDS, 20% (v/v) MeOH) for 25 min at 25 V. The membrane was blocked for at least 1 h with 6% blocking solution (6 g of skim milk powder in 100 mL of Tris-buffered saline with 0.1% (v/v) Tween [TBS-T]). The membrane was washed twice with TBS-T for 10 seconds, after which 3.3 μL of Anti-FLAG® Rb antibody (Sigma-Aldrich) were added to 10 mL of blocking solution and the membrane was left shaking in the solution overnight at 4°C. The membrane was then washed several times with TBS-T, before 3.3 μL of secondary antibody (PierceTM Gt anti-Rb IgG SuperclonalTM secondary antibody HRP conjugate) were added to 10 mL of blocking solution. The membrane was then incubated in the solution for at least 2 h while shaking at room temperature. The membrane was again washed several times with TBS-T, after which proteins were detected using the Bio-Rad Clarity® Western ECL substrate.

### Creation of mutant/overexpression/inducible strains

The in-frame deletion of *rsbN* was created using ReDirect technology. The coding sequence of *rsbN* in cosmid sv-6-D01 was replaced with the *aac(3)IV-oriT* apramycin resistance cassette and the mutant cosmid was confirmed by PCR before being introduced into the non-methylating *E. coli* strain ET12567/pUZ8002 and conjugated into wild type *S. venezuelae.* The in-frame deletion of *chpDFG* and *rdlAB* involved replacing their coding sequences (and intervening genes) in cosmid SV-4-H10 with the *aac(3)IV-oriT* apramycin resistance cassette. As above, the mutant was confirmed with PCR and then conjugated into wild type *S. venezuelae.* Deleting *rnc* from cosmid sv-3-B07 involved replacement with a *hyg-oriT* hygromycin resistance marker. Again, the resulting mutant was confirmed by PCR before being conjugated into the Δ*chpDFG*Δ*rdlAB S. venezuelae* strain to generate the Δ*rnc*Δ*chpDFG*Δ*rdlAB S. venezuelae* strain.

The *phoP* phosphomimetic overexpression plasmid was made by overlap extension-mediated site-directed mutagenesis and subsequent amplification of the resulting *phoP* variant (encoding PhoP D52E). This amplified product was then digested with NdeI and XhoI and cloned into the integrating plasmid pMS82, under the control of the constitutive *ermE\** promoter. The integrity of the resulting construct was confirmed by sequencing prior to conjugation into wild type, *rnc* and *rnj* mutant strains of *S. venezuelae*.

Our thiostrepton-theophylline inducible system was generated by amplifying the *tipA* gene from *Streptomyces coelicolor* genomic DNA and the *tipA* promoter sequence from the pIJ6902 vector. The primers used to amplify these fragments contained the theophylline-responsive riboswitch sequence, such that it was situated between the *tipA* promoter and gene following overlap extension PCR. The overlap extension PCR product (p*tipA-*riboswitch-*tipA*) was digested with BglII and cloned into the integrating plasmid pIJ6902. Constructs with *rnc* and *rnj* fused to an N-terminal 3xFLAG tag were synthesized by GenScript. The sequences for FLAG-tagged *rnc* and *rnj* were PCR amplified, digested with NdeI and KpnI, and cloned behind the original *tipA* promoter in pIJ6902. The resulting constructs had two *tipA* promoters: one driving the expression of *tipA* (with the associated riboswitch) and the other driving the expression of FLAG-tagged *rnc* or *rnj*.

### Transcriptional reporter assays for differentially expressed genes in *rnc* and *rnj* mutants

To test the promoter activity of *glnA* and *nirB,* their promoter sequences were amplified (using primers detailed in Supplementary Table S8), before being digested with XbaI and KpnI and cloned into pGus (Supplementary Tables S7-S8). The resulting constructs were confirmed by sequencing and were introduced into wild type *S. venezuelae* and the RNase mutant strains by conjugation, alongside a promoterless pGUS control. The resulting pGUS-containing strains were inoculated into 10 mL MYM medium and grown overnight. Cultures were normalized to an optical density at 600 nm (OD_600_) of 0.1, and these subcultures were then grown at 30°C for 18 h, after which 1 mL of culture was removed, cells were pelleted and then washed with 1 mL of sterile water.

Washed cell pellets were resuspended in lysis buffer (50 mM phosphate buffer [pH 7.0], 0.27% [v/v] β-mercaptoethanol, 0.1% [v/v] Triton X-100, 1 mg/mL lysozyme) and incubated at 37°C for 30 min. The resulting cell lysate was then centrifuged, and the supernatant was used in the β-glucuronidase assay. In a 96 well plate, 150 μL of Z-buffer (600 mM Na_2_HPO_4_, 400 mM NaH_2_PO_4_, 0.1 mM MgSO_4_, 10 mM KCl, 2.7 mL β-mercaptoethanol), 16.7 μL of cell lysate, and 33.3 μL of PNPG (4 mg/mL p-nitrophenyl-β-D-glucuronide dissolved in Z-buffer) were loaded into each well. For blanks, 166.7 μL Z-buffer and 33.3 μL of PNPG were loaded. OD_420_ was measured at 1 min intervals for 1 h in a microplate reader (CytationTM3 Multi-Mode Microplate Reader and Cell Imager). Miller units were calculated for each replicate at the endpoint (1 h or at time of signal overflow) reading using the following formula:

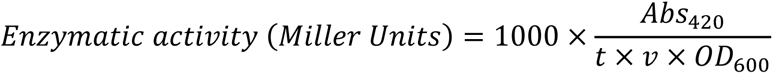

In this calculation, Abs_420_ corresponds to the yellow colour of the o-nitrophenol, v represents volume assayed in milliliters, t represents the reaction time in minutes, and OD_600_ is the optical density at 600 nm of the starting culture and represents cell density. For each of four biological replicates, this measurement was averaged across three technical replicates. The signal associated with the pGUS plasmid control in a respective parent strain (wild type or RNase mutant with/without *phoP*) was subtracted from the experimental *glnA* or *nirB* sample. Subsequently, the fold change between the parent (RNase mutant or wild type) and same strain carrying the *phoP* phosphomimetic overexpression construct was calculated and plotted.

### Chloramphenicol quantification by LC-MS

Cultures were lyophilized and the resulting lyophiles were resuspended in 10 mL methanol and shaken overnight on a rotary shaker at 4°C. After centrifugation to remove particulate matter, the soluble samples were used for LC-MS analyses. The extracts were analyzed using an Agilent 1290 infinity II LC coupled to an Agilent 6495C triple quadrupole, where 5 μL of the injected extracts were separated on a Zorbax SB-C18 column (100 mm by 2.1 mm by 1.8 μm) at a flow rate of 0.3 mL/min. The draw speed was 50 μL/min and the eject speed was 200 μL/min with a 2.4 s wait time. Extracted metabolite separation was achieved using a gradient of 0 to 8 min from 95% to 5% A, 8 to 10 min isocratic 5% A, a gradient of 10 to 10.1 min from 5% to 95% A, and 10.1 to 12 min isocratic 95% A, where A is water with 0.1% formic acid (FA) and B is acetonitrile with 0.1% FA. Chloramphenicol was detected using the negative ionization mode, where 321 – 152 is the quantifying transition and 321 – 35.2 is the qualifying transition (considering precursor ion – product ion). Chloramphenicol was quantified from cell extracts by comparison to a ten-point standard curve of 10 – 2000 ppb chloramphenicol in technical duplicate.

