## Supplementary Tables S6, S7 and S8 for "Defining the networks that connect RNase III and RNase J-mediated regulation of primary and specialized metabolism in *Streptomyces venezuelae*"

**Supplementary Table S6 – Strains used in this study**

| Strains | Genotype/characteristics/use | Reference |
|---|---|---|
| *Streptomyces venezuelae* NRRL B-65442 | Wild type | [1] |
| E325 | *S. venezuelae rnc::aac(3)IV* (*vnz_26040*) | [2] |
| E326 | *S. venezuelae rnj::aac(3)IV* (*vnz_26680*) | [2] |
| E390 | *S. venezuelae chpDFG rdlAB::aac(3)IV* (*vnz_22870-vnz_22985*) *rnc:: hyg* (*vnz_26040*) | This work |
| E391 | *S. venezuelae rsbN::aac(3)IV* (*vnz_15660*) | This work |
| *Streptomyces coelicolor* A3(2) M145 | Wild type; used to amplify *tipA* | [3] |
| *Escherichia coli* DH5α | Routine cloning | Invitrogen |
| *E. coli* BW25113/pIJ790 | Introducing mutations in cosmid DNA | [4] |
| *E. coli* ET12567/pUZ8002 | Generation of methylation-free plasmid DNA and conjugation into *Streptomyces* | [4] |

**Supplementary Table S7 – Plasmids and cosmids used in this study**

| Cosmid/plasmid | Description | Reference |
|---|---|---|
| sv-6-D01 | *S. venezuelae* cosmid carrying *rsbN* | Gift from M. Buttner |
| sv-4-H10 | *S. venezuelae* cosmid carrying *rdlAB chpDFG* | Gift from M. Buttner |
| sv-3-B07 | *S. venezuelae* cosmid carrying *rnc* | Gift from M. Buttner |
| pIJ773 | Plasmid carrying the *aac(3)IV-oriT* cassette | [4] |
| pIJ10700 | Plasmid carrying the *hyg-oriT* cassette | [5] |
| pMS82 | Integrative cloning vector: *hyg*, *oriT*, *int* ΦBT1, *attP* ΦBT1 | [6] |
| pMC390 | $P_{ermE*}$-*vnz_19585* (*phoP* D52E) cloned into pMS82 | This work |
| pIJ6902 | Integrative cloning vector: *aac(3)IV, tsr*, *oriT*, *int* ϕC31, *attP* ϕC31 | [7] |
| pMC388 | pIJ6902::Apra:Hyg::*tipA*; *tipA* sequence from *S. coelicolor* cloned into BglII site of pIJ6902. Apramycin resistance marker replaced with hygromycin resistance marker. | This work |
| pMC387 | pIJ6902::ΔApra:Hyg::riboswitch::*tipA*; theophylline-responsive riboswitch cloned between the *tipA* promoter and *tipA* coding sequence | Riboswitch sequence [8] This work |
| pMC386 | pIJ6902::ΔApra:Hyg::riboswitch::*tipA*::*rnc3×FLAG*; *rnc* sequence with an N-terminal 3×FLAG tag cloned under *tipA* promoter | This work |

| | | |
|---|---|---|
| pMC389 | pIJ6902::ΔApra:Hyg::riboswitch::*tipA*::*rnj3×FLAG*; *rnj* sequence with an N-terminal 3×FLAG tag cloned under *tipA* promoter | This work |
| pGus | Integrative cloning vector: *aac(s)IV*, *aadA*, *oriT*, *int* ϕC31, *attP* ϕC31, *gusA* | [9] |
| pMC391 | Kanamycin resistance gene (*kan*) cloned into pGus | This work |
| pMC392 | *glnA* promoter sequence cloned upstream of promoterless *gusA* in pGus (pMC391) | This work |
| pMC393 | *nirB* promoter sequence cloned upstream of promoterless *gusA* in pGus (pMC391) | This work |

**Supplementary Table S8 – Oligonucleotides used in this study**

| **Name** | **Sequence (5' – 3')*** | **Use** |
|---|---|---|
| rdlB RED F | GGGCGGCGGGGCCGCAACCGAGCGGAGCGTCTTCGCTCA**ATTCCGGGGATCCGTCGACC** | *chpDFG rdlAB* ReDirect cassette |
| chpG RED R | TCAGCCGCCGTAGCCGCCGTCACCCTTGTCGTGACCGTCTG**TAGGCTGGAGCTGCTTC** | *chpDFG rdlAB* ReDirect cassette |
| Rdl int F | CAGGGTTCCCTGAACAAGCC | *chpDFG rdlAB* mutant check |
| Rdl check up R | TGAAGTGACGGCACACGTA | *chpDFG rdlAB* mutant check |
| rsbN RED F | GGGAGTCGACCGTCATGACGAGAGGAGGTGCCGCCAGTG**ATTCCGGGGATCCGTCGACC** | *rsbN* ReDirect cassette |
| rsbN RED R | GGCGCCCCCCGTCCGTGGGGGCGCCCCTTCTTCTTGTCATG**TAGGCTGGAGCTGCTTC** | *rsbN* ReDirect cassette |
| rsbN up F | CACTCTTTCGTGTGGATGCG | *rsbN* mutant check |
| rsbN down R | AGCGAGTCACCGGGAAGCG | *rsbN* mutant check |
| rsbN in R | AGCGAGTCACCGGGAAGCG | *rsbN* mutant check |
| nirB XbaI F | ATATTCTAGAGGCGGGAACGGGTGCG | Cloning *nirB* promoter into pGus |
| nirB KpnI R | ATATGGTACCACCGGGAAGCGTGCGC | Cloning *nirB* promoter into pGus |
| glnA XbaI F | ATATTCTAGACCAAGATCCGAGTGCTTGCC | Cloning *glnA* promoter into pGus |
| glnA KpnI R | ATATGGTACCCCCACTCCTCCTACTCCC | Cloning *glnA* promoter into pGus |
| KanR F | CACGCTGCCGCAAGCACTCAGG | Cloning Kan resistance |
| KanR R | TCAGAAGAACTCGTCAAGAAGGCGA | Cloning Kan resistance |
| TipA BglII F | ATATAGATCTCTGACCGAGGTGGTTCCTCC | Amplify *tipA* with BglII restriction sites |

| | | |
|---|---|---|
| TipA BglII R | ATACAGATCTGTCTCACCAAGACGCTGGTCG | Amplify *tipA* with BglII restriction sites |
| Ribo TipA F | CCAGCATCGTCTTGATGCCCTTGGCAGCACCCTGCTA AGGAGGCAACAAGGTGAGCTACTCCGTGG | Add riboswitch between *tipA* and promoter (overlap extension) |
| Ribo TipA R | AGGGCATCAAGACGATGCTGGTATCACCGGAACCTAT AGTGAGTCGTACTGACGCCGTCCACGC | Add riboswitch between *tipA* and promoter (overlap extension) |
| PhoP F | CATCATCATATGGTGACCCGAGTGCTTGT | Cloning *phoP* into pMS82 |
| PhoP R | CATCATCTCGAGGGAACACATGAAGGGGC | Cloning *phoP* into pMS82 |
| PhoP D52E F | CTCCTCGAGCTGATGC | Site directed mutagenesis of *phoP* (D52E; overlap extension) |
| PhoP D52E R | GCATCAGCTCGAGGAG | Site directed mutagenesis of *phoP* (D52E; overlap extension) |
| 3xF NdeI F | ATATCATATGGACTACAAGGACCACGACGG | Cloning either *rnc* or *rnj* 3xFLAG (binds FLAG sequence) into pIJ6902 |
| rnj R | CATCATGGTACCCCCGTTCTGGCGGAGC | Cloning *rnj* into pIJ6902 |
| rnc R | ATATGGTACCGACAGCGACTCAACC | Cloning *rnc* into pIJ6902 |

*Bold: resistance cassette-specific sequences; underlined: engineered restriction enzyme sites
