## Supplementary figures for "Defining the networks that connect RNase III and RNase J-mediated regulation of primary and specialized metabolism in *Streptomyces venezuelae*"

### Figure S1

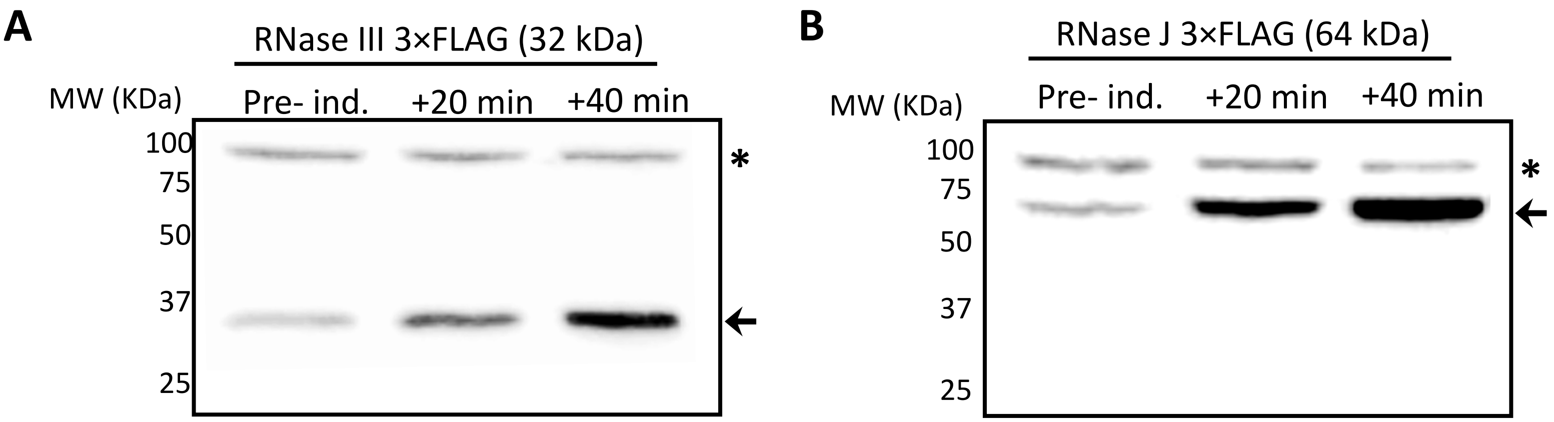


### Figure S2

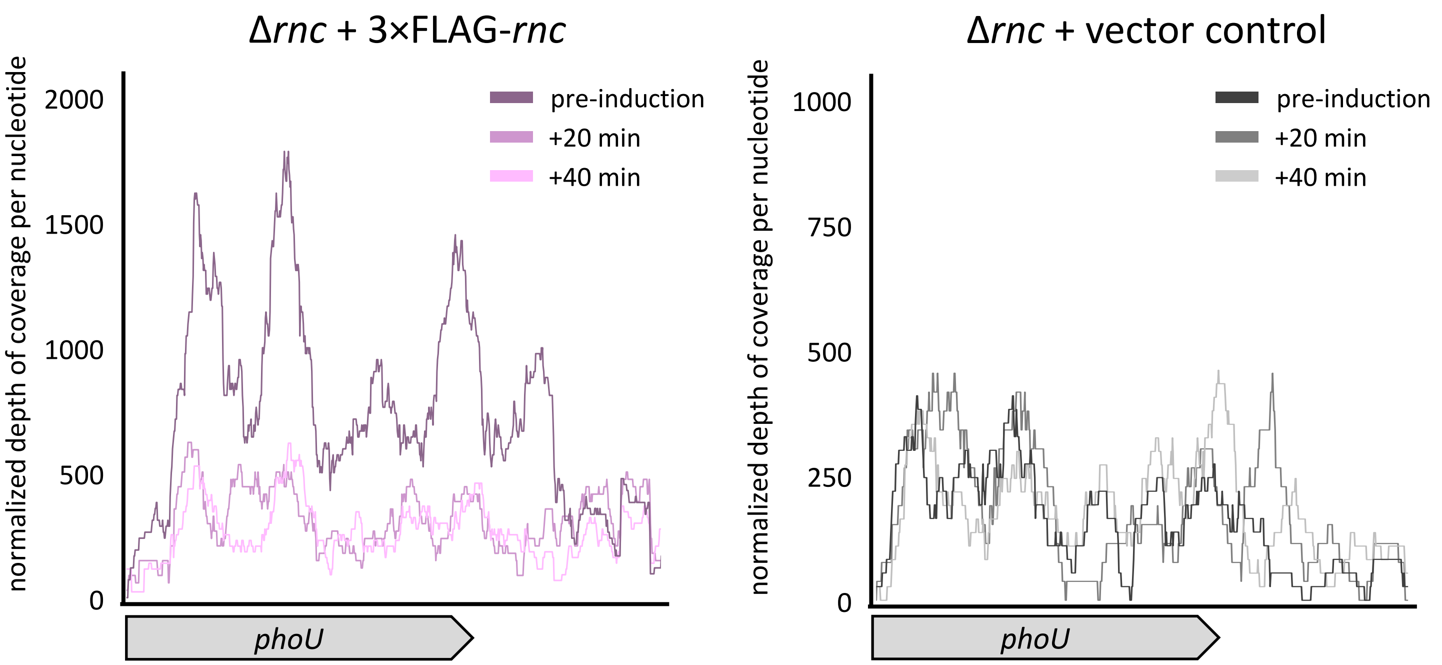


### Figure S3

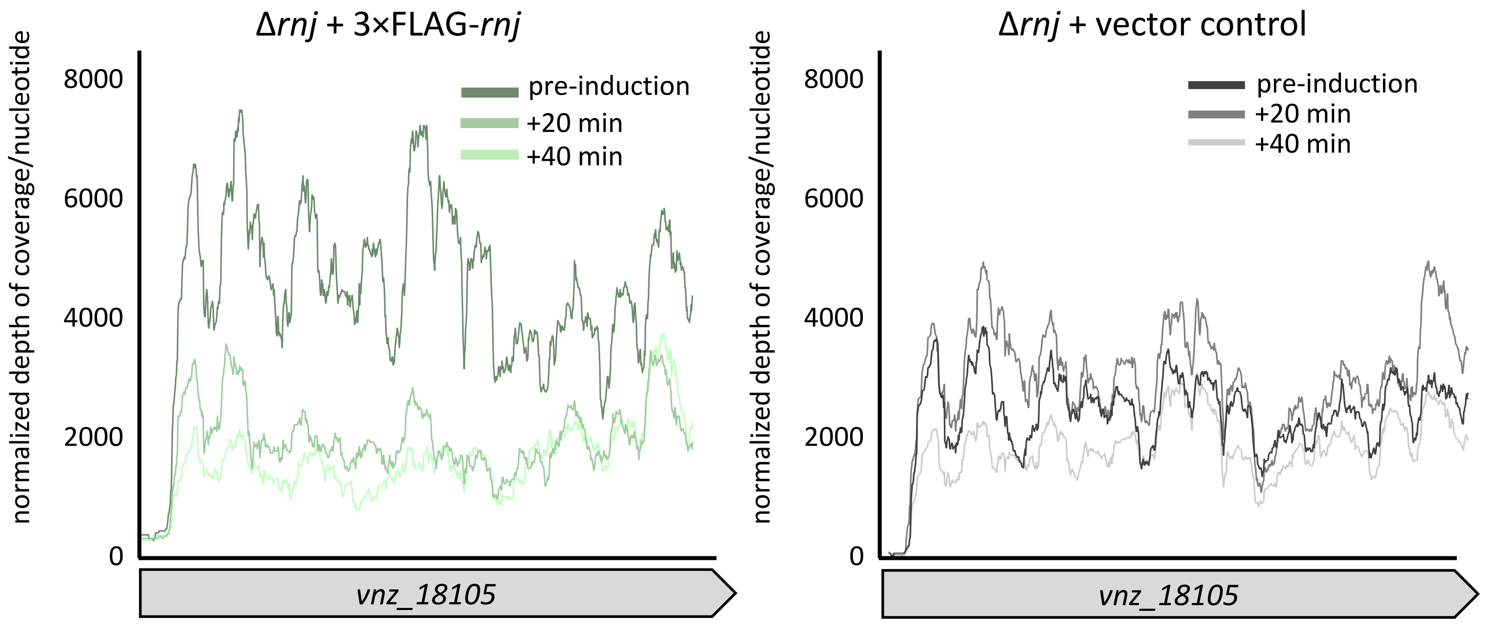

Figure S1

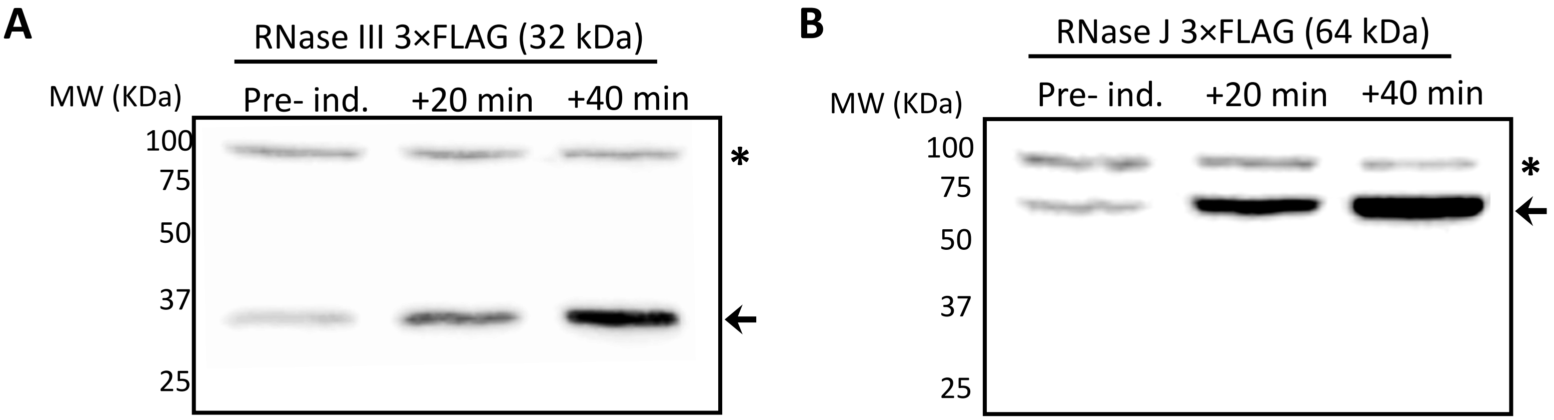


**Supplementary Figure 1: RNase III and RNase J are effectively induced following addition of thiostrepton.** Immunoblot for **(A)** RNase III 3×FLAG and **(B)** RNase J 3×FLAG using anti-FLAG-specific antibodies, in samples isolated pre-induction and 20- and 40-minutes post-induction. Arrow heads indicate the induced RNase III (left) and RNase J (right). Asterisks mark a non-specific band that served as a protein loading control.

### Figure S2

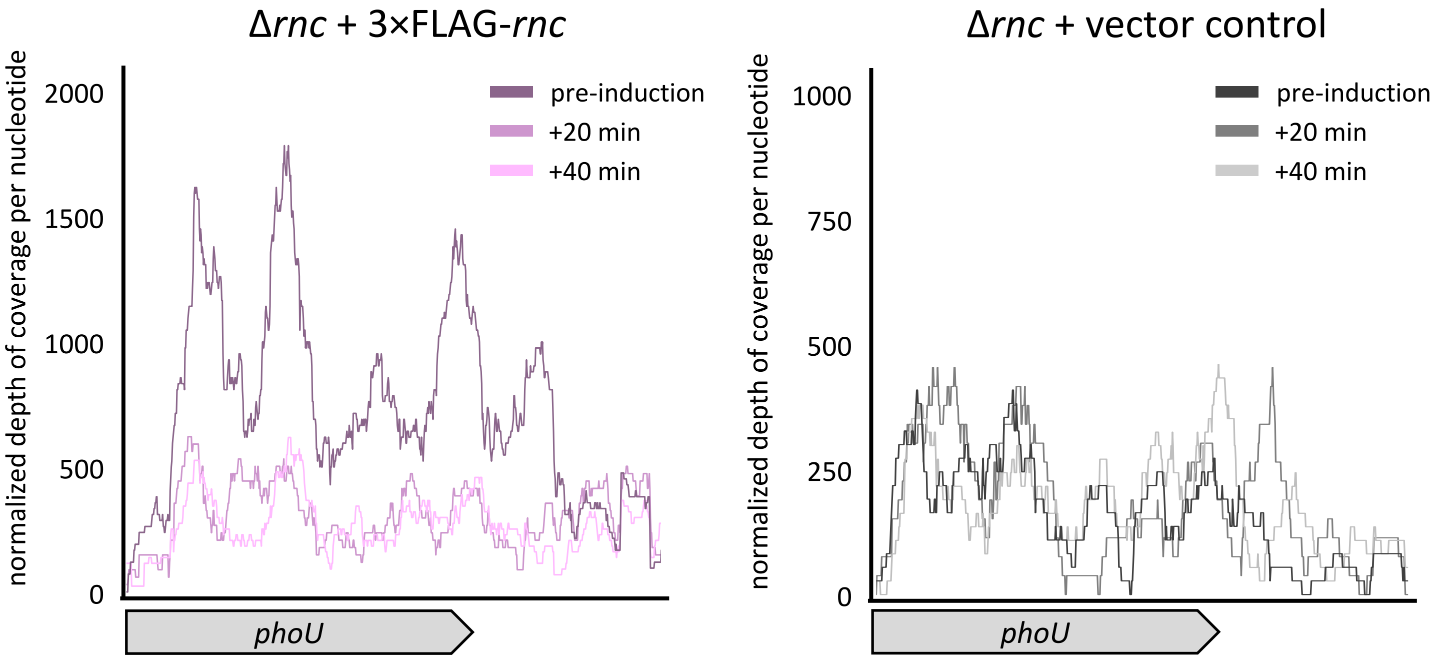


**Supplementary Figure 2: *phoU* transcript is directly targeted by RNase III.** Second biological replicate for experiment presented in Figure 5C.

### Figure S3

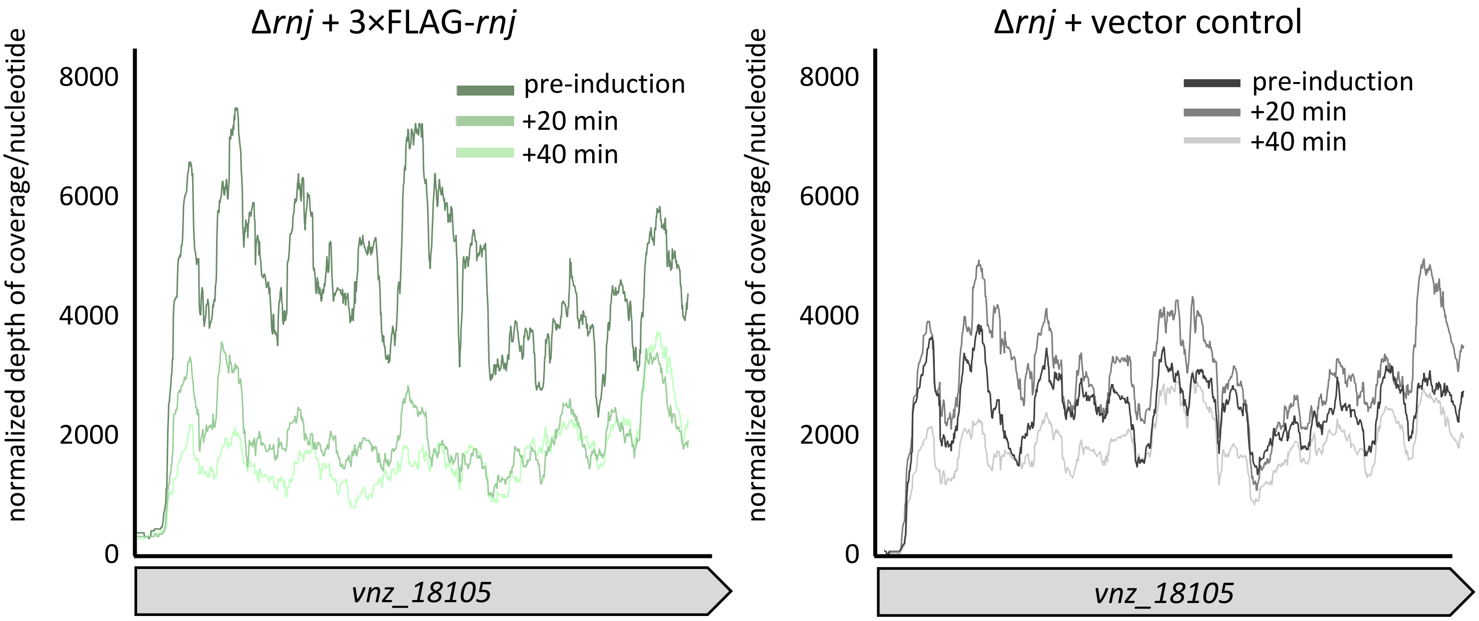


**Supplementary Figure 3: *vnz_18105* transcript is directly targeted by RNase J.** Second biological replicate for experiment presented in Figure 6.

Figure S4

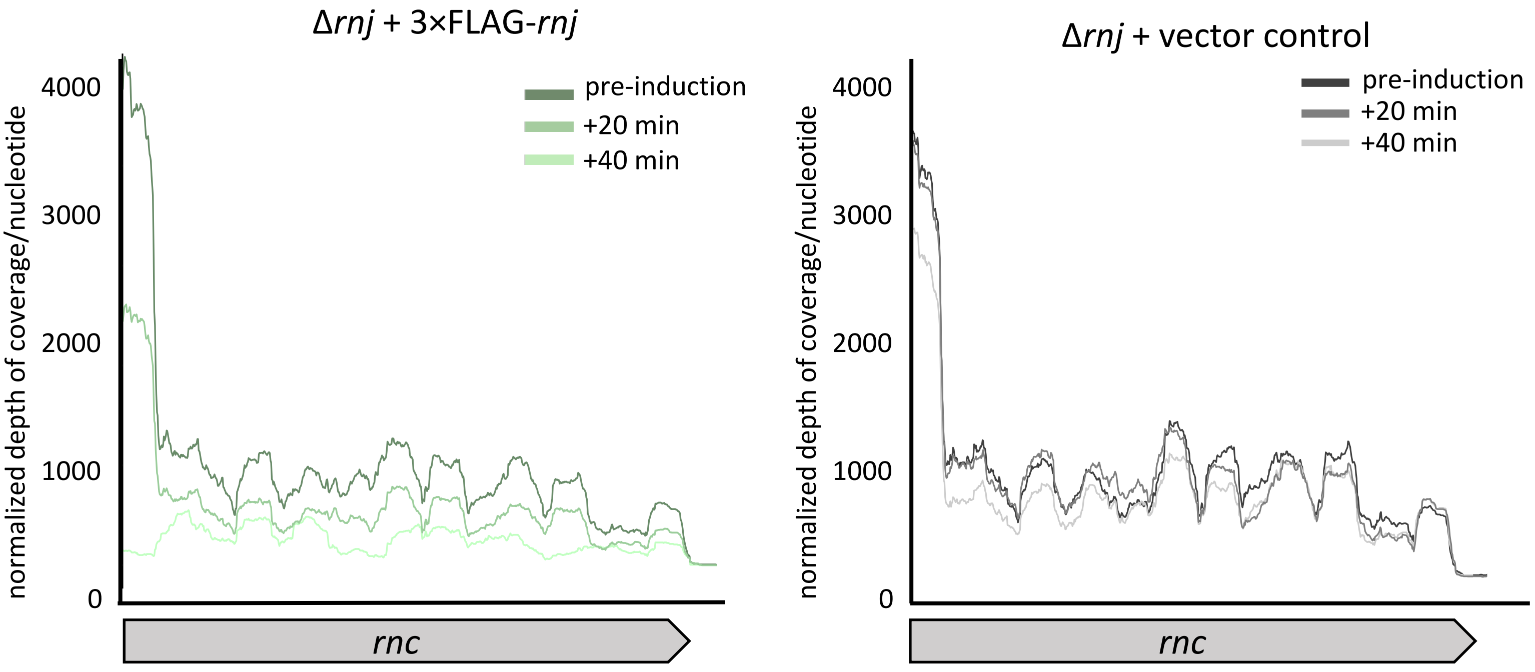


**Supplementary Figure 4: *rnc* transcript is directly targeted by RNase J.** Second biological replicate for experiment presented in Figure 7B.

Figure S5

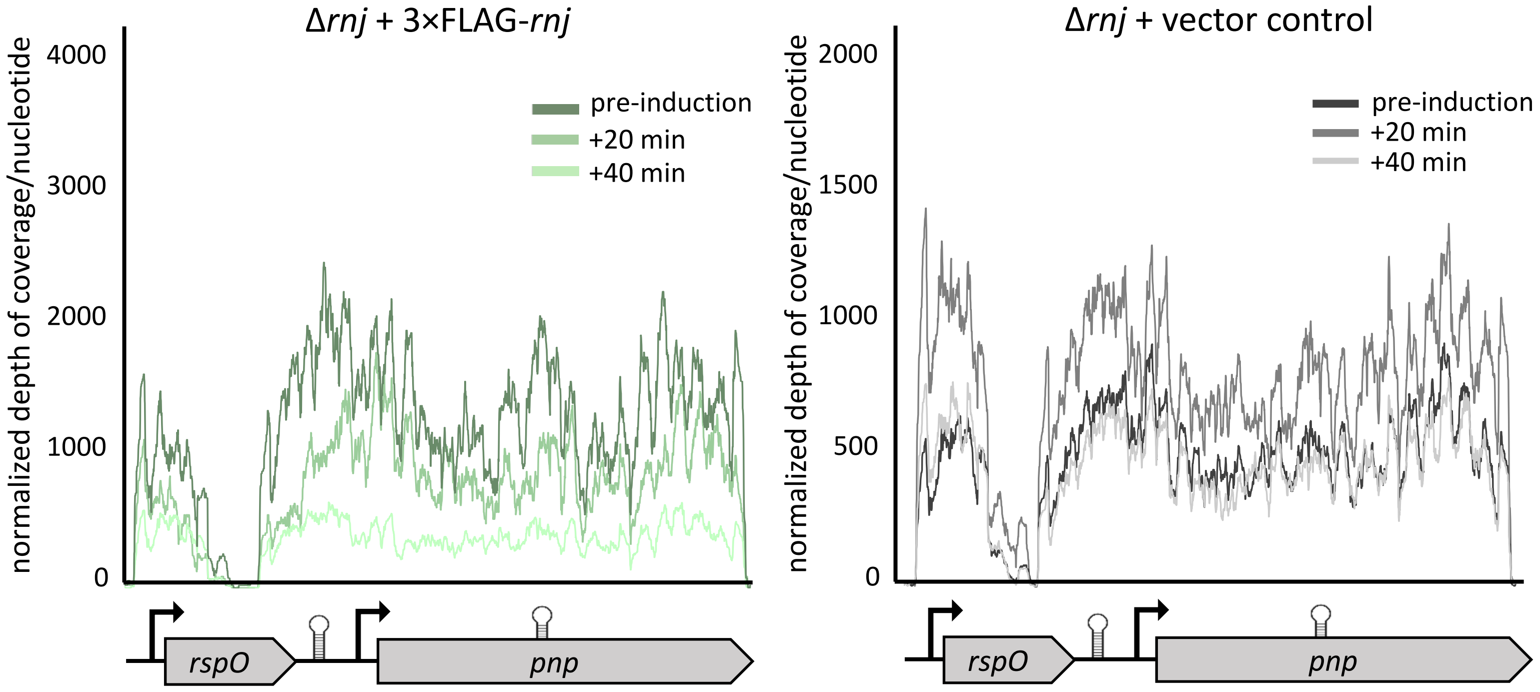


**Supplementary Figure 5: *rspO-pnp* transcript is directly targeted by RNase J.** Second biological replicate for experiment presented in Figure 7D.
